# Synchronized Latency Reversal and Immune Clearance by a Multifunctional Fusion Protein Enables HIV-1 Reservoir Reduction

**DOI:** 10.64898/2025.12.10.693434

**Authors:** Fei Luo, Yinghui Cao, Na Liu, Xiaowei Wang, Wenjia Hu, Xiaoqing Xie, Dongmei Jiang, Yangyang Xu, Jun Wang, Hongyu Wang, Zhenyang Yu, Peng Qian, Shicheng Wan, Zhida Liu, Tielong Chen, Shihui Song, Yong Xiong, Xiaoping Tang, Linghua Li, Li Wang, Haisheng Yu, Liang Cheng

## Abstract

Strategies that concurrently reactivate latent reservoirs and enhance immune-mediated clearance hold significant promise for achieving an HIV cure. Here, we developed hyperIL-15×sCD4-Fc (15×sCD4-Fc), a fusion protein that integrates latency reactivation, targeted immune engagement, and effector-mediated killing into a single molecule. This agent not only potently reverses HIV-1 latency in CD4^+^ T cells from people living with HIV-1 (PLWH) through coordinated IL-15 receptor signaling and sCD4-mediated HIV-1 envelope (Env) engagement, but also enhances antigen-specific CD8^+^ T cell response in PBMCs derived from PLWH. Furthermore, 15×sCD4-Fc enables Env-specific elimination of reactivated latently infected cells by NK cells while preventing off-target cytotoxicity. In PBMCs from PLWH, 15×sCD4-Fc reduced replication-competent HIV-1 DNA by 93.8%. In antiretroviral treated HIV-1-infected humanized mice, the molecule demonstrated both safety and efficacy in diminishing viral reservoir in lymphoid organs. This spatiotemporally coupled approach to reservoir exposure and immune recognition establishes a clinically viable strategy for clearing the HIV-1 reservoir.

**Graphical abstract:** 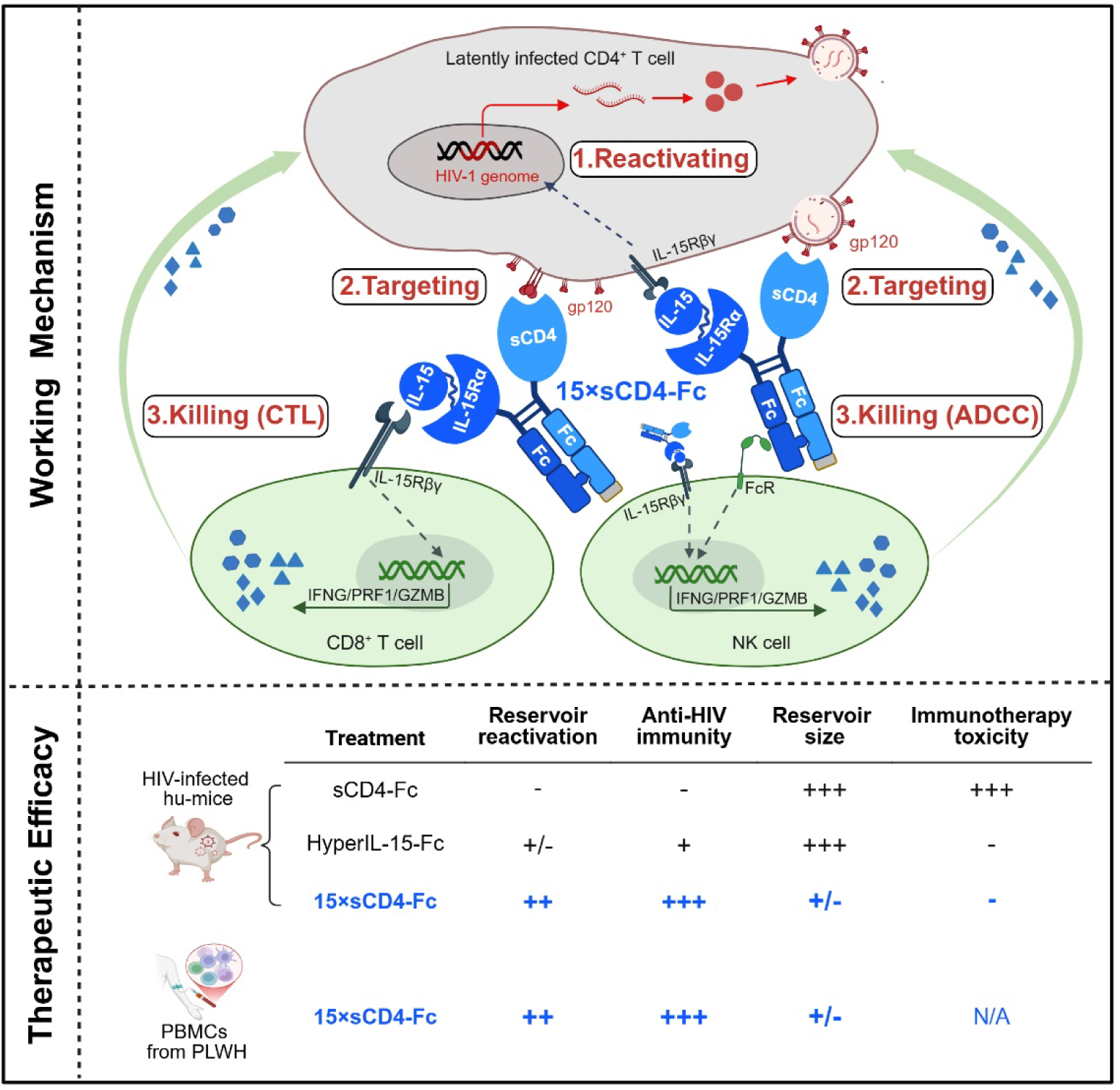

## INTRODUCTION

Despite the remarkable success of antiretroviral therapy (ART) in suppressing HIV-1 replication, the persistent latent viral reservoirs remain the primary obstacle to achieving a cure.^1,2^ These transcriptionally silent reservoirs harbor intact proviral DNA that evades immune surveillance while retaining the capacity to drive viral rebound upon treatment interruption.^3–5^ Current eradication strategies hypothesize a three-phase requirement: (i) latency reversal or reactivation, (ii) immune recognition of virus-producing cells, and (iii) effector-mediated elimination.^6,7^ While the “shock-and-kill” paradigm^8,9^ has dominated cure strategies, with latency-reversing agents (LRAs) deployed to expose latent viruses for immune-mediated or viral cytopathic elimination,^5,10–14^ clinical trials consistently show LRA-induced reservoir reactivation fails to achieve meaningful reservoir reduction,^10,11,14–17^ even with adjunct immunotherapies like HIV vaccines^18–20^ or neutralizing antibodies^21–23^ designed to enhance cell clearance. Inadequate spatiotemporal coordination between latency reversal, immune recognition, and targeted elimination likely represents a key driver of this limitation.^7^

We hypothesized that a molecular agent capable of synchronizing viral reservoir activation, immune effector engagement, and targeted elimination could overcome fundamental barriers to HIV-1 eradication. In the present study, we engineered hyperIL-15×sCD4-Fc (15×sCD4-Fc), a triple-armed fusion protein comprising three functional modules: (i) the hyperIL-15 component, a covalent IL-15/IL-15Rα sushi domain complex functioning as an IL-15 superagonist,^24^ selected based on its clinical-stage analog N-803 (IL-15/IL-15Rα-Fc complex) that induces latency reversal in IL-2Rβγ^+^ CD4^+^ T cells^25,26^ and sensitizes reactivated latent reservoirs for CD8^+^ T cell recognition,^25^ while concurrently activating cytotoxic CD8^+^ T cells and NK cells,^27–29^ with these immunostimulatory properties supported by preclinical SIV/HIV-1 models,^30–34^ and clinical trials demonstrating IL-15 superagonist safety in people living with HIV (PLWH);^35^ (ii) the soluble CD4 (sCD4) module, derived from HIV-1’s cellular receptor,^36,37^ which neutralizes free virions^38^ while exposing occluded Env epitopes to enhance antibody-dependent cellular cytotoxicity (ADCC) against infected cells,^39,40^ achieving dual targeting and neutralization; and (iii) the Fc domain that bridges Env^+^ target cells with FcγR^+^ effectors (e.g., NK cells), which have been shown to be critical for the in vivo activity of broadly neutralizing anti-HIV-1 antibodies,^41^ establishing an immunologically active microenvironment for eliminating reactivated reservoirs.

Using primary CD4^+^ T cells from PLWH and an HIV-1 latently infected cell line, we confirmed the individual functionalities of the fusion protein components. More importantly, we discovered synergistic interactions among the three functional domains. The 15×sCD4-Fc mediates synergistic latency reversal through coordinated IL-15 receptor signaling and sCD4-Env engagement. Furthermore, 15×sCD4-Fc establishes a self-contained therapeutic circuit by directly coupling latency reversal with sCD4-Env-targeted immune attack via Fc-dependent NK cell engagement, a mechanism unattainable by hyperIL-15-Fc or its analog N803 alone. In addition, we demonstrated that heterodimeric architecture exploits monomeric sCD4’s lack of binding to MHC-II to ensure HIV-specific targeting without off-target toxicity. Preclinical validation in humanized mouse models and PLWH-derived cells demonstrated that 15×sCD4-Fc surpasses hyperIL-15-Fc or sCD4-Fc monotherapy in: (i) activating latent HIV-1 reservoirs, (ii) enhancing antigen-specific CD8^+^ T cell and NK cell responses, and (iii) reducing intact proviral DNA and replication-competent viral reservoir size. These findings collectively advance the “shock-and-kill” paradigm by engineering a therapeutic framework that overcomes critical barriers in HIV-1 reservoir clearance, offering a translationally viable path toward viral eradication.

## RESULTS

### Development and characterization of fusion protein hyperIL-15×sCD4-Fc

We engineered the multifunctional fusion protein hyperIL-15×sCD4-Fc (15×sCD4-Fc) by integrating three functional modules: (1) an IL-15 superagonist (hyperIL-15) comprising human IL-15 fused to the sushi domain of the IL-15 receptor α chain, as previously reported in our studies,^24,42^ (2) a soluble CD4 (sCD4) module containing the extracellular D1D2 domain of human CD4, and (3) a knob-into-hole modified human IgG1 Fc domain enabling heterodimer formation **(Figure 1A; Figure S1A)**. Non-reducing SDS-PAGE analysis confirmed successful 15×sCD4-Fc heterodimer formation, showing a predominant band at the expected position **(Figure 1B)**. Under reducing conditions, the dissociated constituent hyperIL-15-Fc6 and sCD4-Fc9 showed expected molecular weights **(Figure S1B)**. Corresponding homodimeric controls (hyperIL-15-Fc and sCD4-Fc) were generated for functional comparisons **(Figure S1C-H)**.

**Figure 1.**
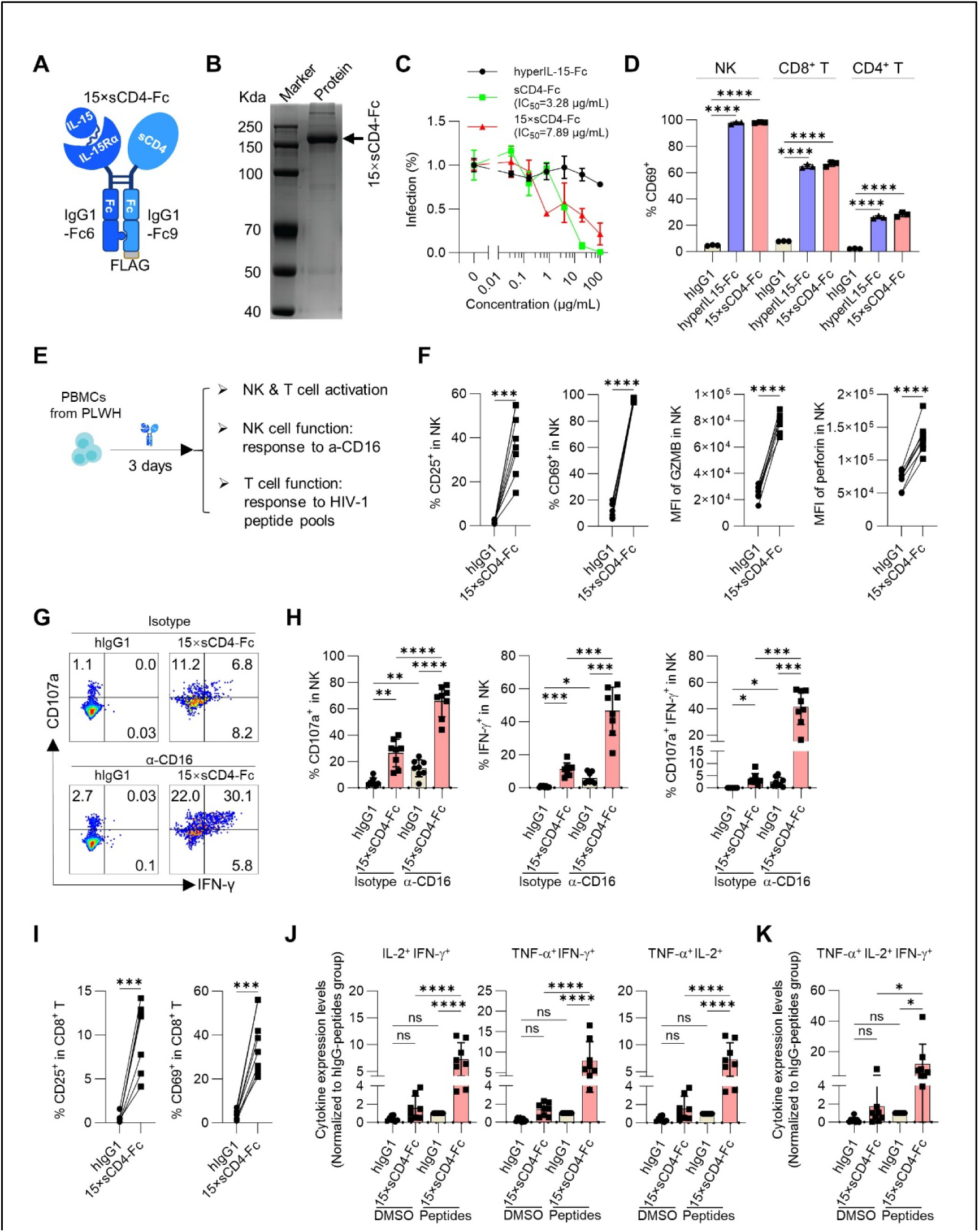
Development and characterization of multifunctional fusion protein hyperIL-15×sCD4-Fc. **(A)** Schematic of the 15×sCD4-Fc heterodimic fusion protein architecture. **(B)** SDS-PAGE (non-reducing condition) of purified 15×sCD4-Fc, showing a predominant band. **(C)** Neutralizing cell-free HIV-1. HIV-1_JRCSF_ virions were pre-incubated with serially diluted proteins (100∼0.032 μg/mL) before infecting TZM-bl cells. Infectivity was quantified by luciferase activity. **(D)** Activation of human NK and T cells. PBMCs from healthy donors (n=3) were treated with indicated proteins (10 nM) for 48 hours, followed by CD69 detection via flow cytometry. Data represent mean ± SEM of triplicate measurements from one representative donor. **(E)** Schematic of experimental design. PBMCs from ART-suppressed PLWH (n=8 donors) were treated with 15×sCD4-Fc or hIgG1 control in the presence of efavirenz. The activation and function of NK cells and T cells were analyzed after 3 days of culture. **(F)** NK cells activation. Summary data show expression of CD25, CD69, granzyme B (GZMB) and perforin on NK cells at day 3. **(G-H)** NK cells activity. On day 3, PBMCs were stimulated with α-CD16 antibody or isotype control for 6 hours (Brefeldin A added at 1 hour). Representative flow cytometry plots (G) and summary data (H) show CD107a and IFN-γ expression in NK cells. **(I)** CD8^+^ T cells activation. Summary data show expression of CD25 and CD69 on CD8^+^ T cell at day 3. **(J-K)** Antigen-specific CD8^+^ T cell responses. On day 3, PBMCs were stimulated with HIV-1 Env/Gag/Pol peptide pools for 8 hours (Brefeldin A added at 3 hours). Summary data show intracellular IFN-γ, IL-2 and TNF-α double-positive (J) and triple-positive (K) CD8^+^ T cells detected by flow cytometry. Data in (D, H, J and K) are presented as mean ± SEM. Data in (F and I) are presented as paired data plot. Statistical significance was determined by one-way ANOVA with Tukey’s post hoc test (D, H, J and K) or paired Student’s t-test (F and I). \**p* < 0.05, \*\**p* < 0.01, \*\*\**p* < 0.001, \*\*\*\**p* < 0.0001.

First, we test the activity of the engineered sCD4 and hyperIL-15 domains of the fusion protein. Neutralization assays against HIV-1_JRCSF_ demonstrated that while homodimeric sCD4-Fc exhibited an IC50 of 3.28 μg/mL (66.0 nM sCD4 molecules), the heterodimeric 15×sCD4-Fc maintained comparable neutralization potency per sCD4 domain (IC50 = 7.89 μg/mL; 79.4 nM sCD4 molecules) **(Figure 1C)**. Notably, a 50% reduction in sCD4 valency only modestly affected neutralization efficiency, with a 1.2-fold difference in per-domain activity, indicating preserved Env binding function despite the structural modifications. As control, hyperIL-15-Fc showed no neutralization activity **(Figure 1C)**. Using PBMCs from healthy donors, NK and T cell activation assays revealed that both 15×sCD4-Fc and hyperIL-15-Fc efficiently activated NK cells, CD8^+^ T cells, and CD4^+^ T cells, as indicated by CD69 upregulation **(Figure 1D)**. These results confirm that the sCD4 and hyperIL-15 domains retain their respective HIV-1-neutralizing and immunostimulatory function within the fusion protein.

To further determine whether 15×sCD4-Fc enhances NK cell cytotoxicity and HIV-specific T cell responses from people living with HIV (PLWH), we cultured the protein with PBMCs isolated from ART-suppressed PLWH **(Figure 1E, Table S1)**. Compared to the hIgG1 control, 15×sCD4-Fc significantly activated NK cells from PLWH, as indicated by upregulated expression of CD25 and CD69 **(Figure 1F)**. NK cells from 15×sCD4-Fc-treated PBMCs also exhibited enhanced cytotoxic potential, demonstrated by increased intracellular expression of granzyme B and perforin **(Figure 1F)**. Following restimulation with α-CD16 antibody, NK cells from 15×sCD4-Fc-treated group showed elevated surface CD107a expression and higher IFN-γ production **(Figures 1G and 1)**. 15×sCD4-Fc treatment also activated CD8^+^ T cells from PLWH (**Figure 1I**). Upon HIV-1 peptide restimulation, 15×sCD4-Fc-treated samples exhibited superior polyfunctional CD8^+^ T cell responses, with higher frequencies of cells double-positive or triple-positive for IFN-γ, IL-2, and TNF-α **(Figure 1J and 1; Figures S2A and S2B)**. Minimal cytokine production was detected in unstimulated controls (15×sCD4-Fc without peptides) or in peptide-stimulated hIgG1 samples **(Figure 1J and 1; Figures S2A and S2B)**, confirming the antigen-specific nature of the response. Although 15×sCD4-Fc activated CD4^+^ T cells from PLWH **(Figure S2C)**, cytokine production by these cells following re-stimulation with HIV-1 peptide pools showed minimal change compared to vehicle control **(Figures S2D and S2E)**, suggesting antigen-nonspecific CD4^+^ T cells activation.

Collectively, these results demonstrate the successful design and functional validation of the heterodimeric fusion protein 15×sCD4-Fc.

### 15×sCD4-Fc outperforms hyperIL-15-Fc in reversing HIV-1 latency

To evaluate the latency-reversing capacity of 15×sCD4-Fc, we utilized primary CD4^+^ T cells from ART-suppressed PLWH who had maintained a plasma viral load below 20 copies/mL for over 12 months (**Table S1**). After isolating high-purity CD4^+^ T cells (>99%; **Figure S3A**), we treated them with 15×sCD4-Fc, hyperIL-15-Fc, sCD4-Fc, or hIgG1 control in the presence of antiretrovirals (elvitegravir plus efavirenz) for 72 hours and then quantifying HIV-1 RNA in supernatant **(Figure 2A; Figure S3B)**.

**Figure 2.**
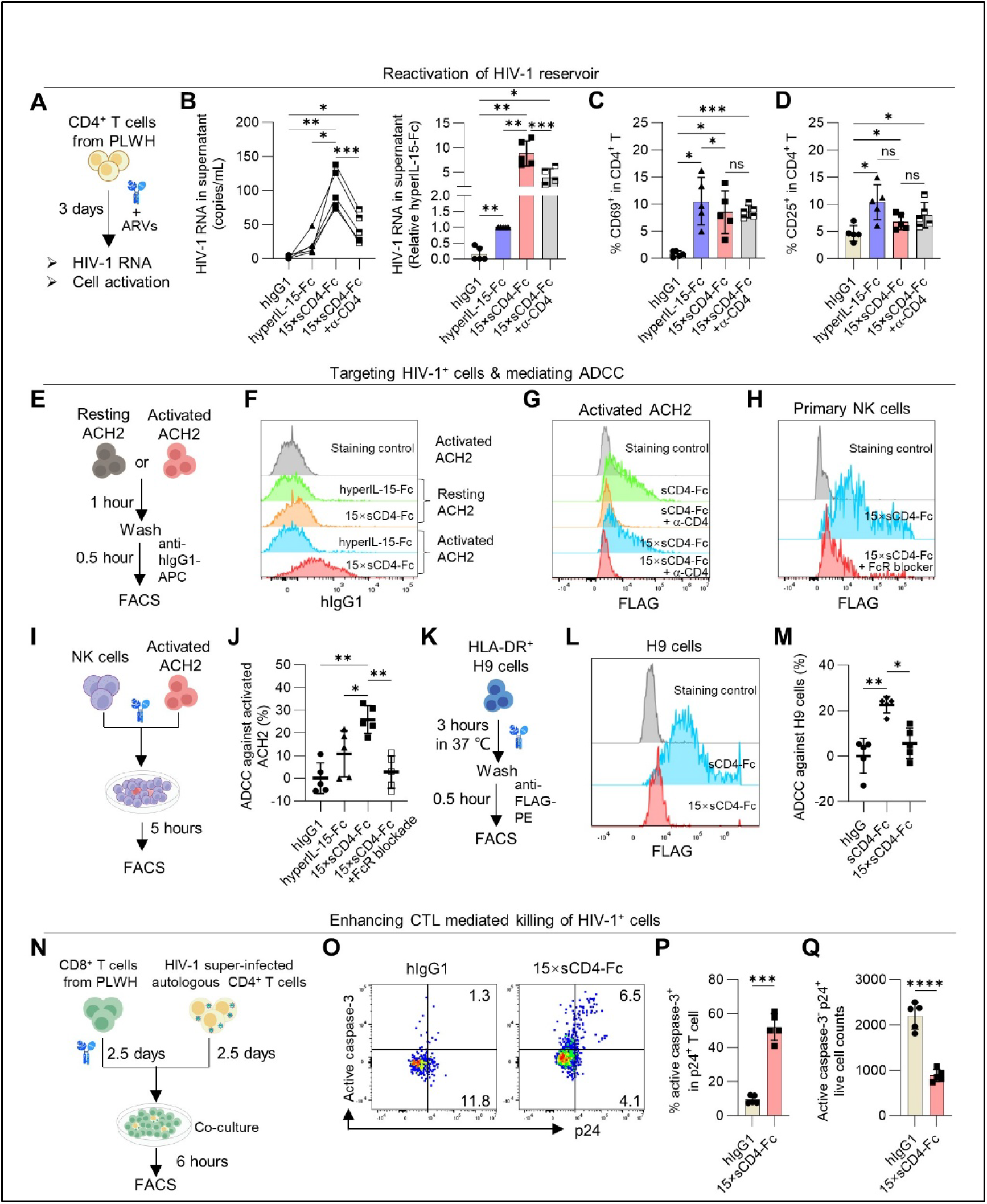
15×sCD4-Fc reactivates HIV-1 reservoirs, targets infected cells, and mediates elimination via ADCC and CTL activity. **(A-D)** Reservoir reactivation capacity. (A) Experimental schematic: primary CD4⁺ cells isolated from cART-suppressed subjects (n=5 donors) were cultured for 72 hours with fusion proteins (10 nM) in ARVs-containing media, followed by HIV-1 detection in supernatant and flow cytometric analysis of cells activation. For blocking the interaction between HIV-1 gp120 and sCD4, α-CD4 monoclonal antibody (clone RPA-T4, 50 nM) was pre-incubated with 15×sCD4-Fc (10 nM) in room temperature for 20 minutes and then added to CD4^+^ T cells. (B) Absolute (left) and normalized (right) HIV-1 RNA levels in supernatant. (C-D) Summary data show expression of CD69 (C) and CD25 (D) on CD4^+^ T cells. **(E-G)** Binding specificity to HIV^+^ cell. (E) Experimental schematic: PMA-activated or resting ACH2 cells were incubated with 20 nM hyperIL-15-Fc or 15×sCD4-Fc, followed by anti-human IgG1-APC staining. For blocking the interaction between HIV-1 gp120 and sCD4, α-CD4 monoclonal antibody (clone RPA-T4, 50 nM) was pre-incubated with sCD4-Fc or 15×sCD4-Fc in room temperature for 20 minutes and then incubated with activated ACH2 cells. (F-G) Representative flow histograms (n=2 experiments). **(H)** Fc receptor binding capacity. PBMCs from healthy donors were pre-treated with or without FcR blockade followed by incubation with 15×sCD4-Fc (20 nM) for 30 minutes, and staining with anti-FLAG, CD3 and CD56. Representative flow histograms showed the binding of 15×sCD4-Fc to CD3^-^CD56^+^ NK cells. **(I-J)** ADCC-mediated HIV-1^+^ cell killing. (I) Experimental schematic: primary human NK cells (effectors) were co-cultured with CFES-labeled (5 µM) PMA-activated ACH2 cells (targets) or CFES-labeled (0.5 µM) non-activated ACH2 cells (control) at a 30:1 effector-to-target (E:T) ratio for 5 hours, followed by viable cell quantification via flow cytometry. (J) Specific lysis percentages were calculated. **(K-M)** Binding and killing of HLA-DR^+^ H9 cells. (K) Experimental schematic: H9 cells were incubated with 20 nM sCD4-Fc or 15×sCD4-Fc for 3 hours in 37 ℃, followed by anti-FLAG-APC staining. (L) Representative flow histograms (n=2 experiments). (M) Primary human NK cells (effectors) were co-cultured for 6 hours at 30:1 E:T ratio with H9 cells (targets) that had been pre-treated for 3 hours with indicated proteins. Viable H9 cells were quantified by flow cytometry. Specific lysis percentages were calculated. **(N-Q)** CTL-mediated elimination. (N) Experimental schematic: CD4^+^ T cells (targets) from cART-suppressed subjects were superinfected with HIV_JR-CSF_ and co-cultured for 6 hours with autologous CD8^+^ T cells (effectors; 20:1 E:T ratio) that had been pre-treated with 15×sCD4-Fc. Viable cells were quantified by flow cytometry. (O, P) Representation flow plots (O) and summarized data (P) show percentage of active caspase-3^+^p24^+^ populations in CD4^+^ T cells. (Q) Absolute number of viable caspase-3^-^ p24^+^ CD4^+^ T cells. Data in (B right, C, D, J, M, P and Q) are presented as mean ± SEM. Data in (B left) are presented as paired data plot. Data in (J, M, P and Q) are representative of one experiment, presented as mean ± SEM from two independent experiments. Statistical significance was determined by one-way ANOVA with Tukey’s post hoc test (B-D, J and M) or unpaired Student’s t-test (P and Q). \**p* < 0.05, \*\**p* < 0.01, \*\*\**p* < 0.001, \*\*\*\**p* < 0.0001.

Consistent with previous reports on N803,^25,26^ hyperIL-15-Fc induced a 6.5-fold increase in viral production **(Figure 2B)**, which correlated with its CD4^+^ T cell activation potential as measured by CD69 and CD25 upregulation **(Figures 2C and 2)**. Remarkably, 15×sCD4-Fc achieved significantly enhanced proviral reactivation (57.5-fold increase), substantially exceeding the effect of hyperIL-15-Fc alone **(Figure 2B)**. This enhancement was particularly notable given that 15×sCD4-Fc contains only half the hyperIL-15 content of homodimeric hyperIL-15-Fc and showed similar CD4^+^ T cell activation **(Figures 2C and 2**; **Figure 1D)**. To elucidate the mechanistic basis of this synergy, we employed an anti-CD4 antibody (clone RPA-T4) which targets the D1-domain of sCD4 and competitively inhibits gp120 binding^43^. Antibody-mediated disruption of sCD4-Env interaction reduced 15×sCD4-Fc’s latency-reversing activity by 56.3%, while maintaining its CD4^+^ T-cell activation capacity **(Figure 2B-D)**, directly implicating Env engagement in the observed synergy. As control, sCD4-Fc showed no detectable latency-reversing activity **(Figure S3C and S3D)**. We propose a model whereby sCD4 binding to hyperIL-15-induced Env retains 15×sCD4-Fc on reactivated reservoir cells, thereby enabling sustained IL-15 signaling and consequently enhanced viral production.

These results suggest that 15×sCD4-Fc’s engineered dual-module architecture mediates superior latency reversal via coordinated IL-15 receptor signaling and Env engagement, providing a promising strategy for HIV-1 reservoir targeting.

### 15×sCD4-Fc specifically targets reactivated reservoir cells for ADCC while sparing MHC-II-expressing cells

We next investigated whether 15×sCD4-Fc could specifically target reactivated HIV-1-infected cells and mediating antibody-dependent cellular cytotoxicity (ADCC). Using the well-characterized ACH2 latency model where PMA stimulation induces HIV-1 antigen expression,^44^ we performed flow cytometric analysis comparing binding to activated versus resting cells **(Figure 2E)**. 15×sCD4-Fc showed selective binding to PMA-activated cells but not their resting counterparts, while hyperIL-15-Fc control exhibited no detectable binding regardless of activation status **(Figure 2F)**. The binding of 15×sCD4-Fc or sCD4-Fc to activated ACH2 can be blocked by the anti-CD4 antibody (clone RPA-T4) **(Figure 2G)**. This activation-dependent binding pattern confirms the sCD4 module’s ability to specifically recognize Env-expressing reactivated reservoir cells. In addition, flow cytometric analysis confirmed Fc-dependent binding of 15×sCD4-Fc to human primary NK cells and monocytes **(Figure 2H; Figure S4A)**, suggesting preserved Fc functionality critical for effector cell engagement. Functional NK cell co-culture assays demonstrated that 15×sCD4-Fc potently induced ADCC against reactivated ACH2 cells **(Figure 2I and 2)**. This effect was completely abolished by Fc receptor blockade **(Figure 2J)**, confirming Fc-dependent mechanisms. This ADCC activity was specific to 15×sCD4-Fc, as the hyperIL-15-Fc control exhibited minimal cytotoxicity **(Figure 2J)**.

We next addressed potential safety concerns stemming from CD4’s dual role as both an HIV-1 receptor and a T cell coreceptor for MHC class II (MHC-II) interactions.^36,37,45,46^ Using HLA-DR-expressing H9 T cells **(Figure S4B)**, we observed strong binding with homodimeric sCD4-Fc but negligible interaction with heterodimeric 15×sCD4-Fc **(Figure 2K and 2)**. Consistent with these findings, 15×sCD4-Fc showed no ADCC activity against H9 cells in coculture with NK cells, whereas homodimeric sCD4-Fc triggered ADCC **(Figure 2M)**. The MHC-II-dependent nature of this effect was confirmed through competitive inhibition using DR-12a, an HLA-DR β1-derived peptide,^47^ which reduced sCD4-Fc binding to H9 cells **(Figure S4C)** and completely blocked ADCC activity **(Figure S4D)**. Comparable inhibition was achieved through Fc mutations (P329G/L234A/L235A; FcMut) that abrogate Fc receptor binding^48^ **(Figure S4D)**.

Taken together, these results demonstrated that 15×sCD4-Fc specifically targets reactivated reservoir cells for ADCC while sparing MHC-II-expressing cells.

### 15×sCD4-Fc enhances CD8^+^ T cell-mediated clearance of autologous HIV-1-infected CD4^+^ T cells from PLWH

Our data demonstrate that 15×sCD4-Fc enhances antigen-specific CD8^+^ T cell response in PBMCs from PLWH **(Figure 1I-K)**. To determine whether 15×sCD4-Fc could facilitate CD8^+^ T cell-mediated clearance HIV-1^+^ CD4^+^ T cells, we established autologous co-cultures using CD8^+^ T cells and HIV-1-infected CD4^+^ T cells from ART-suppressed individuals **(Figure 2N)**. In superinfected CD4^+^ T cell co-cultures, pre-exposure of CD8^+^ T cells with 15×sCD4-Fc increased the percentage of active caspase-3^+^ p24^+^ target CD4^+^ T cells by 5.6-fold **(Figure 2O and P)** and reduced live p24^+^ cell numbers by 60% **(Figure 2Q)**, confirming direct cytotoxic T lymphocytes (CTL)-mediated killing of reactivated reservoirs.

Collectively, these results demonstrate that 15×sCD4-Fc uniquely integrates three essential therapeutic functions within a single molecule: (i) reservoir reactivation via IL-15R signaling and enhanced by sCD4, (ii) precision targeting through sCD4-Env interaction, and (iii) immune-mediated clearance via both NK cell ADCC and CTL cytotoxicity **(Figure 3A)**. This multifunctional architecture addresses critical limitations of current latency-reversing approaches by coupling viral reactivation with targeted immune effector engagement.

**Figure 3.**
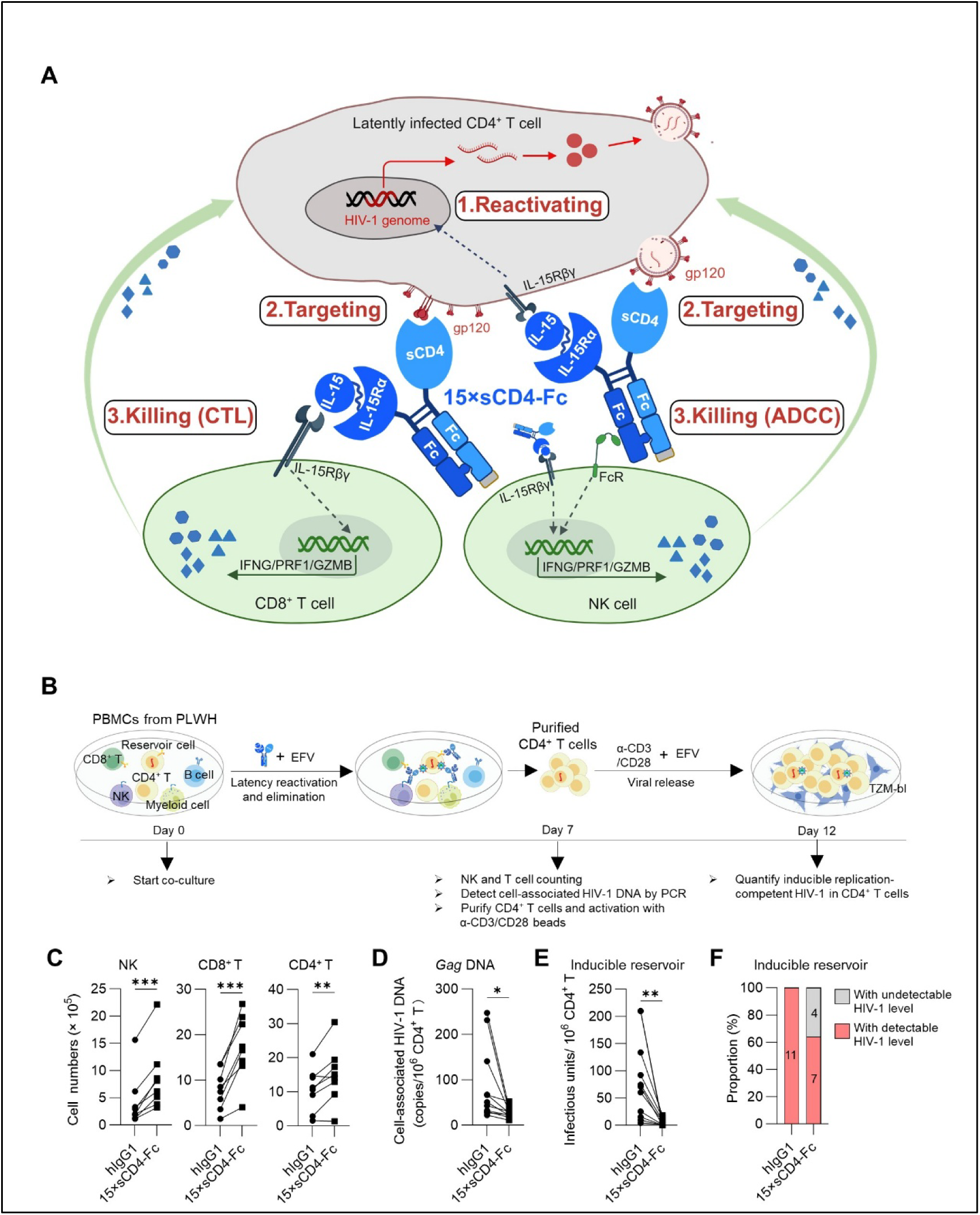
15×sCD4-Fc treatment reduces HIV-1 reservoirs in ART-suppressed PBMCs from PLWH. **(A)** The proposed working model illustrates how 15×sCD4-Fc coordinates three synergistic actions to overcome the spatiotemporal barrier in HIV-1 cure: it first reactivates the latent reservoir via hyperIL-15 signaling, inducing viral antigen expression; the sCD4 domain then specifically targets these reactivated cells by binding to newly surfaced Env; finally, the molecule enhances the function of HIV-1-specific CD8^+^ T cells and NK cells, orchestrating targeted clearance through CTL and ADCC mechanisms. This schematic was created with BioRender (https://BioRender.com/e7o3n31). **(B)** Schematic of experimental design. PBMCs from ART-suppressed PLWH were treated with 15×sCD4-Fc or hIgG1 control in the presence of efavirenz for 7 days. Live CD4^+^ T cells were sorted, stimulated with α-CD3/CD28 beads (efavirenz-maintained) for 5 days, and quantified for replication-competent HIV-1 using TZM-bl assay. **(C)** NK cell and T cell expansion. Summary data show absolute numbers of NK cells, CD8^+^ T cells, and CD4^+^ T cells on day 7 (n=8 donors). **(D)** Cell-associated HIV-1 DNA reduction. HIV-1 *gag* DNA in purified CD4^+^ T cells at day 7 of culture was quantified by qPCR (n=11 donors). **(E)** Inducible replication-competent HIV-1 reservoir reduction. Replication-competent HIV-1 viruses in CD4^+^ T cells were detected by TZM-bl assay (n=11 donors). **(F)** Frequency of samples with inducible viruses. Data in (C-E) are presented as paired data plot. Statistical significance was determined by paired Student’s t-test (C-E). \**p* < 0.05, \*\**p* < 0.01, \*\*\**p* < 0.001.

### 15×sCD4-Fc treatment reduces HIV-1 reservoirs in ART-suppressed PBMCs from PLWH

To evaluate the therapeutic potential 15×sCD4-Fc in reducing HIV-1 reservoirs, we established an ex vivo system assess reservoir eradication **(Figure 3B)**. PBMCs from ART-suppressed PLWH **(Table S1)** were treated with 15×sCD4-Fc or hIgG1 control in the presence of efavirenz for 7 days. On day 7, live CD4^+^ T cells were sorted, and cell-associated HIV-1 DNA was detected by PCR. To measure replication-competent HIV-1, the sorted CD4^+^ T cells were also stimulated with α-CD3/CD28 beads for 5 days (with continued efavirenz treatment), and viral output was quantified using ultrasensitive TZM-bl assays.^49^

After 7 days of treatment, 15×sCD4-Fc induced 2.0-fold expansion of NK cells, 2.2-fold expansion of CD8^+^ T cells, and 1.4-fold expansion of CD4^+^ T cells **(Figure 3C)**. Quantitative PCR analysis demonstrated a 68.2% reduction in cell-associated HIV-1 *gag* DNA in purified CD4^+^ T cells compared to hIgG1-treated controls **(Figure 3D)**. Strikingly, quantification of replication-competent HIV-1 revealed a 93.8% decrease in infectious units per million (IUPM) CD4^+^ T cells, from 82.3 ± 70.6 IUPM in controls to 5.1 ± 6.2 IUPM in treated samples (**Figure 3E**). Furthermore, 36% (4/11) of 15×sCD4-Fc-treated samples showed suppression of viral production below the detection limit in TZM-bl assays, compared to 0% in control groups **(Figure 3F)**.

These results demonstrate that 15×sCD4-Fc enhances HIV-1-specific immunity while substantially reducing replication-competent reservoirs in patient-derived cells, highlighting its therapeutic potential for HIV-1 cure strategies.

### 15×sCD4-Fc eliminates MHC-II-dependent bystander cytotoxicity associated with sCD4-Fc in HIV-1-infected humanized mice

Previous clinical studies reported that administration of CD4-IgG1 or CD4-IgG2 (PRO 542) at doses up to 3 mg/kg in subjects with AIDS was well tolerated, with no significant clinical or immunological toxicities and no anti-CD4-IgG antibody induction.^54–56^ However, a re-analysis of data from one study using CD4-IgG1, which included detailed CD4^+^ T cell count data^54^ revealed divergent outcomes in immunological toxicities when participants were stratified into two groups based on study completion **(Figure S5)**. Among 22 HIV-1-positive participants, the 14 subjects who completed the 12-week treatment exhibited a significant 37.4% decrease in CD4^+^ T cell counts, whereas the 8 participants who discontinued treatment prematurely showed no such reduction **(Figure S5)**. These results suggest potential immunotoxicity of CD4-IgG1 in humans.

To assess the immunological safety of 15×sCD4-Fc in vivo, we utilized our well-established humanized mouse model of HIV-1 infection.^50,51^ Humanized mice (hu-mice) with HIV-1 infection received combination antiretroviral therapy (cART) at week 3 post-infection and were administered four doses of 15×sCD4-Fc, sCD4-Fc, or sCD4-FcMut twice weekly starting from week 7 **(Figure 4A; Table S2)**. As expected, cART effectively suppressed viremia in all animals within 4 weeks **(Figure 4B)**. While neither sCD4-Fc nor sCD4-FcMut treatment altered HIV-1 viremia during cART, transient viral blips were detected in 15×sCD4-Fc-treated mice **(Figure 4B)**, suggesting reservoir reactivation by 15×sCD4-Fc.

**Figure 4.**
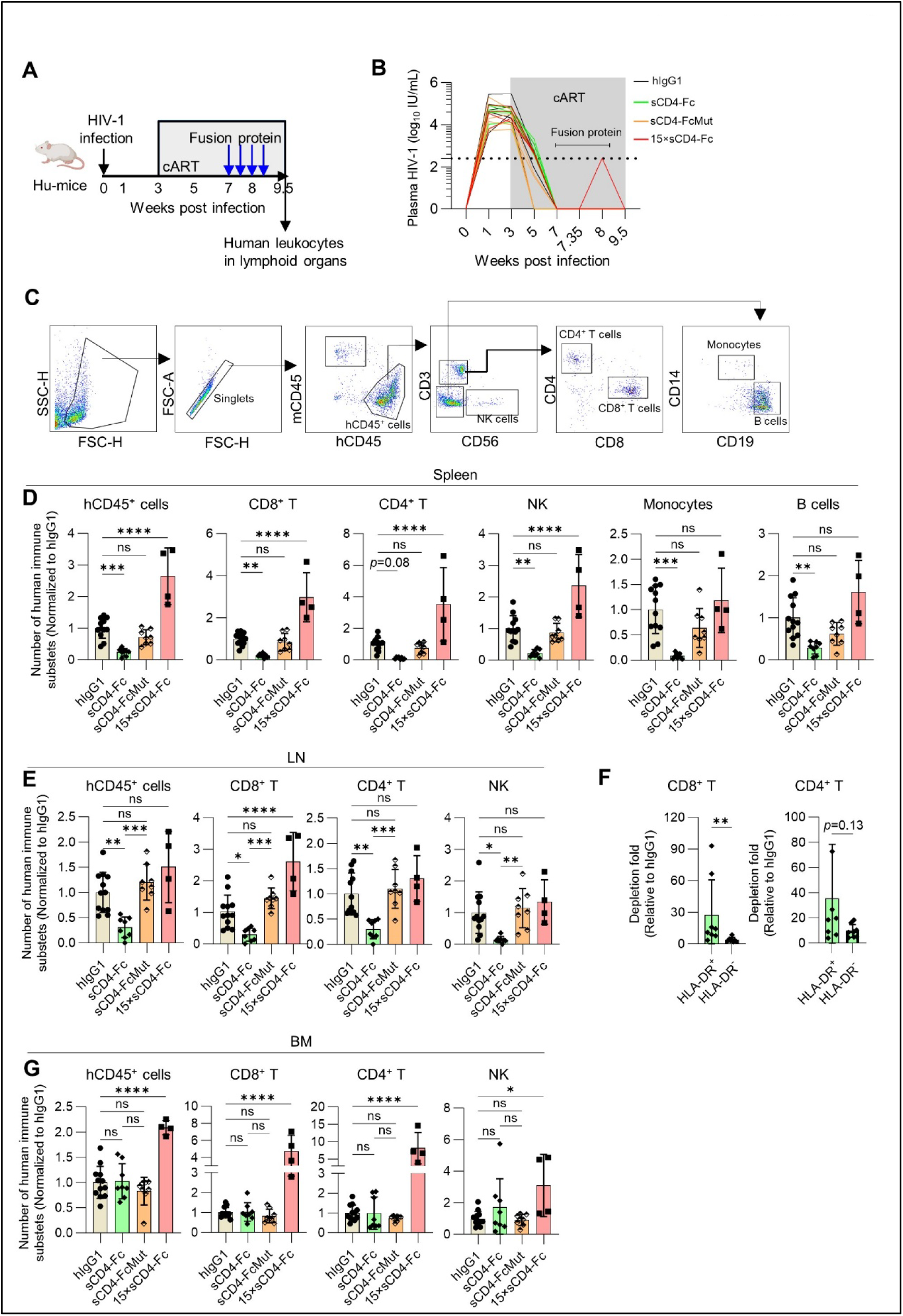
15×sCD4-Fc eliminates MHC-II-dependent bystander cytotoxicity associated with sCD4-Fc in HIV-1-infected humanized mice. **(A)** Schematic diagram of in vivo experiment. HIV-1-infected hu-mice received cART from 3-9.5 weeks post-infection (wpi), with test proteins (hIgG1/sCD4-Fc/sCD4-FcMut/15×sCD4-Fc; 20 μg/mouse) administered twice weekly during 7-9 wpi (hIgG1 (n=12), sCD4-Fc (n=8), sCD4-Fc-FcMut (n=8), 15×sCD4-Fc (n=4); combined from 2 independent experiments). HIV-1 RNA in plasma of hu-mice quantified by RT-qPCR at indicated timepoints. Human CD45^+^ leukocytes numbers in tissues were counted. **(B)** HIV-1 RNA levels in the plasma. Dashed line indicates assay limit of detection. **(C)** Gating strategy for detecting human immune subsets in spleens of hu-mice by flow cytometry. **(D)** Relative numbers of total hCD45^+^ leukocytes, human CD8^+^ T cells, CD4^+^ T cells, NK cells, monocytes, and B cells in spleens at endpoint. **(E)** Relative numbers of total hCD45^+^ leukocytes, human CD8^+^ T cells, CD4^+^ T cells and NK cells in lymph nodes at endpoint. **(F)** The relative depletion of HLA-DR^+^ versus HLA-DR^-^ T cell subpopulations in spleens of sCD4-Fc-treated groups compared to hIgG1-treated controls, expressed as fold reduction. **(G)** Relative numbers of total hCD45^+^ leukocytes, human CD8^+^ T cells, CD4^+^ T cells and NK cells in bone marrow at endpoint. Data in (D-G) are presented as mean ± SEM. Statistical significance was determined by one-way ANOVA followed by Tukey’s multiple comparisons test (D, E and G) or unpaired Student’s t-test (F). \**p* < 0.05, \*\**p* < 0.01, \*\*\**p* < 0.001, \*\*\*\**p* < 0.0001.

Consistent with its ability to bind to HLA-DR^+^ cells and mediating ADCC in vitro, administration of the control protein sCD4-Fc induced systemic depletion of human CD45^+^ leukocytes, including CD8^+^ T cells, CD4^+^ T cells, NK cells, monocytes, and B cells, in both splenic **(Figure 4C and 4)** and lymph node compartments **(Figure 4E)**. In marked contrast, 15×sCD4-Fc not only avoided leukocyte depletion but significantly expanded human T-cell and NK cell populations **(Figure 4D and 4)**. The immunotoxicity of sCD4-Fc preferentially affected HLA-DR^+^ T cells **(Figure 4F)** and was entirely abolished by Fc inactivation (sCD4-FcMut; **Figure 4D and 4)**, confirming MHC-II- and Fc-dependent mechanisms. Supporting this observation, sCD4-Fc-mediated toxicity was absent in the bone marrow **(Figure 4G)**, a known sanctuary site resistant to antibody-mediated clearance.^52,53^

Collectively, these results demonstrate that whereas the homodimeric sCD4-Fc induces Fc-dependent, MHC-II-driven bystander cytotoxicity, the heterodimeric 15×sCD4-Fc maintains stringent Env specificity and expands antiviral NK and CD8^+^ T cells, thereby achieving efficacy without off-target toxicity.

### 15×sCD4-Fc achieves tripartite HIV-1 reservoir control through coordinated reactivation, immune potentiation and viral clearance in humanized mice

Building upon its multifunctional and safety properties, we systematically evaluated the therapeutic potential of 15×sCD4-Fc in cART-suppressed, HIV-1-infected humanized mice. Treatment regimens (15×sCD4-Fc, hyperIL-15-Fc, or hIgG1 control) were initiated at 7 weeks post-infection, following the achievement of cART-mediated viral suppression, and administered for 2-3 weeks. Serum viremia was monitored throughout the treatment period, while immune profiles and lymphoid reservoirs were quantified at 1 week after treatment cessation **(Figure 5A; Table S3)**.

**Figure 5.**
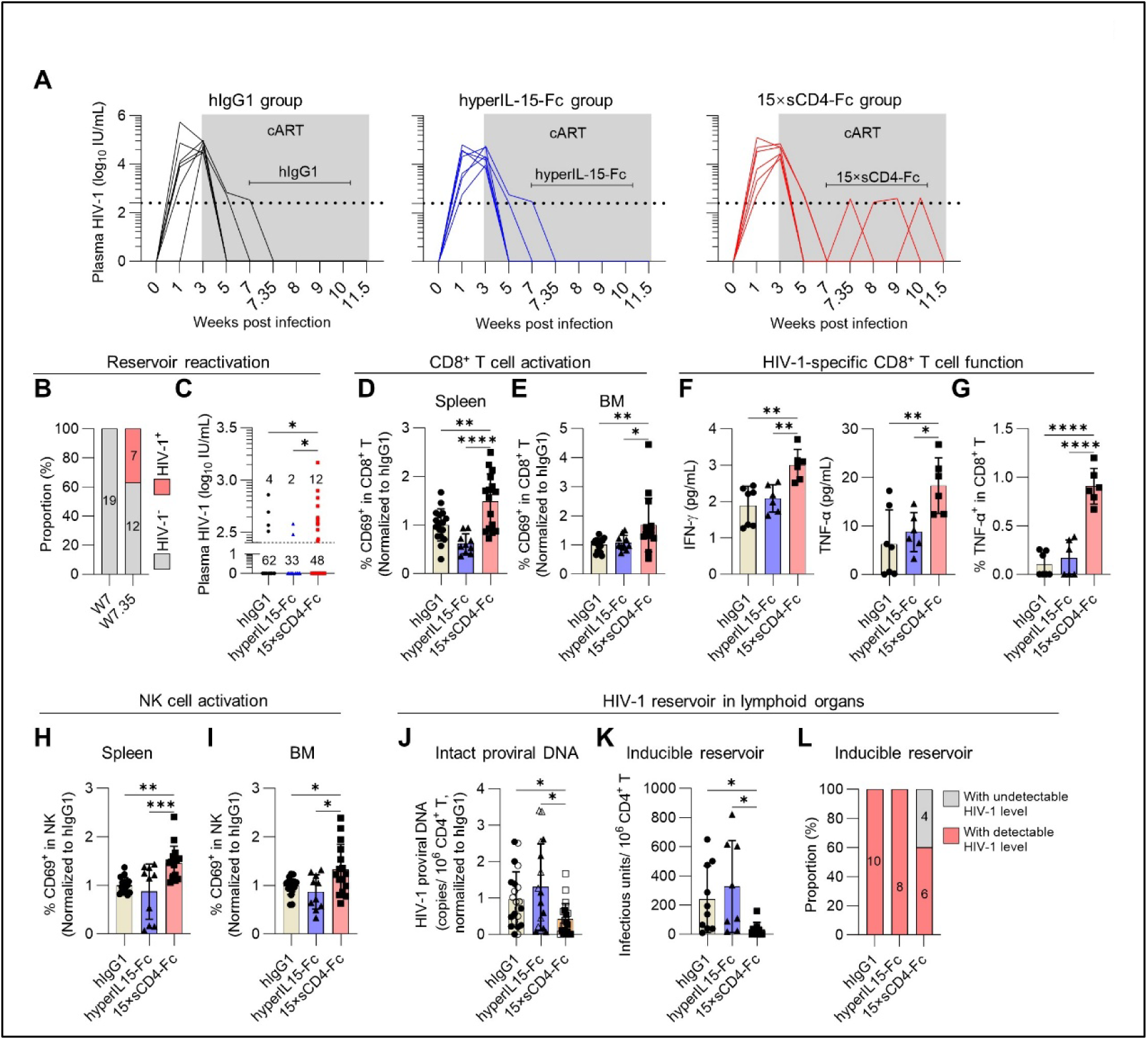
15×sCD4-Fc reactivates HIV-1 reservoirs, enhances antiviral immunity, and reduces reservoir size in hu-mice. HIV-1-infected hu-mice initiated cART at 3 wpi. Beginning at 7 wpi, cART-suppressed mice received twice-weekly intraperitoneal injections of hIgG, hyperIL-15-Fc or 15×sCD4-Fc (20 µg/mouse) for 2-3 weeks. All animals were euthanized 1 week after the final treatment administration. **(A-C)** Viral suppression and reactivation kinetics. (A) Plasma HIV-1 RNA levels in a representative cohort of infected hu-mice following treatment. Dashed line indicates assay limit of detection. (B) Percentage of hu-mice with detectable viremia at 60 hours post-injection of 15×sCD4-Fc across 5 independent cohorts. (C) Pooled analysis of viremia at multiple time points (weeks 7.35, 8, 9, and 10) after injection of hIgG1, hyperIL-15-Fc, and 15×sCD4-Fc across five independent cohorts. Numbers above the dash line in C indicate the number of samples with detectable viremia; numbers below indicate samples without detectable viremia. **(D-E)** CD8^+^ T cell activation. Relative CD69 expression on CD8^+^ T cells from spleens (D) and bone marrow (E). **(F-G)** Antigen-specific CD8^+^ T cell responses. Mixture of splenocytes and BM cells from hu-mice were stimulated ex vivo with HIV-1 Env/Gag/Pol peptide pools in the presence of α-CD28 and CD49d. (F) IFN-γ and TNF-α secretion in supernatant after 24 hours were detected by ELISA. (G) Intracellular TNF-α in CD8^+^ T cells following 8-hour peptide stimulation (Brefeldin A added at 3 hours) were detected by flow cytometry. **(H-I)** NK cell activation. Relative CD69 expression on NK cells from spleens (H) and bone marrow (I). **(J)** Reservoir quantification by IPDA. Pooled data show intact proviral DNA levels in spleens (open symbols) and BM (solid symbols) (hIgG1, n=11; hyperIL-15-Fc, n=8; 15×sCD4-Fc, n=11, Combined data from 4 independent experiments). **(K-L)** Inducible replication-competent reservoir analysis. (K) Infectious units per million (IUPM) CD4^+^ T cells in BM detected by TZM-bl assay. (L) Frequency of samples with inducible virus (hIgG1, n=10; hyperIL-15-Fc, n=8; 15×sCD4-Fc, n=10; Combined data from 4 independent experiments). Data in (D-K) are presented as mean ± SEM. Data in (B-E, H and I) represent combined results from 5 independent mouse cohorts (hIgG1, n=17; hyperIL-15-Fc, n=10; 15×sCD4-Fc, n=16). Statistical significance was determined by one-way ANOVA followed by Tukey’s post hoc multiple comparisons test (D, E, H, I) or non-parametric Kruskal-Wallis *H* test (J and K). \**p* < 0.05, \*\**p* < 0.01, \*\*\**p* < 0.001, \*\*\*\**p* < 0.0001.

Extensive preclinical studies have established that N-803 (a clinical-stage IL-15 superagonist analog of hyperIL-15-Fc), while capable of latent reservoir reactivation in vivo, requires concomitant CD8^+^ T cell depletion to achieve measurable viral rebound in ART-treated SIV-infected macaques and HIV-1-infected humanized mice.^32,33^ Mirroring this phenotype, hyperIL-15-Fc therapy in our system likewise failed to induce detectable reservoir reactivation **(Figure 5A-C; Table S3)**. In marked contrast, 15×sCD4-Fc treatment elicited viremia blips during treatment **(Figure 5A-C and 4B; Tables S2-3)**, with quantifiable plasma HIV-1 RNA detected in 7 out of 19 animals (37%) at 60 hours post-treatment **(Figure 5B)** and 12 out of 60 samples (20%) collected at all timepoints **(Figure 5C)**. This robust HIV-1 reservoir reactivation profile highlights 15×sCD4-Fc’s unique capacity to bypass the immunological constraints that limit current IL-15 or N803-based latency reversal strategies.

Comprehensive immunological characterization demonstrated the superior immune-stimulatory capacity of 15×sCD4-Fc compared to hyperIL-15-Fc. Both treatments inducing CD8^+^ T cell expansion in splenic and bone marrow compartments, yet only 15×sCD4-Fc achieved statistically significant increases **(Figure S6A)**. Notably, 15×sCD4-Fc-treated mice exhibited uniquely elevated expression of the activation marker CD69 on CD8^+^ T cells relative to all control groups **(Figure 5D and 5)**, while functional assessment revealed markedly enhanced cytotoxic potential characterized by significantly increased production of granzyme B and interferon-γ following either PMA/ionomycin treatment **(Figure S6B)** or anti-CD3/CD28 stimulation **(Figure S6C)**. Enhanced antigen-specific responses were evidenced by elevated IFN-γ and TNF-α secretion along with increased frequency of TNF-α^+^ CD8^+^ T cells following HIV-1 peptide pool stimulation **(Figure 5F and 5)**. In contrast, hyperIL-15-Fc failed to enhance any measured parameter of CD8^+^ T cell functionality across all assays **(Figure 5F and 5; Figure S6B and S6C)**. This result is consistent with published primate studies demonstrating that combined N-803 and broadly neutralizing antibodies (bNAbs) treatment, but not N-803 alone, enhanced the multifunctionality of antiviral CD8^+^ T cells in SHIV-infected, ART-suppressed macaques.^57^ Although both treatments increased human NK cells in spleen and bone marrow **(Figure S6D; Figure 4D and 4)**, only 15×sCD4-Fc treatment significantly activated human NK cells in both compartments, with hyperIL-15-Fc showing no comparable effect **(Figure 5H and I)**. Collectively, these results underscore the unique immunological advantages of 15×sCD4-Fc for HIV-1 reservoir control in vivo.

Reservoir reduction analysis demonstrated HIV-1 reservoir depletion following 15×sCD4-Fc treatment. The Intact Proviral DNA Assay (IPDA) confirmed a 68.1% decrease in intact proviral DNA loads **(Figure 5J)**, while TZM-bl assay (TZA), which quantifying inducible replication-competent reservoirs through sensitive reporter cell detection,^49^ showed a particularly robust 86.2% reduction in bone marrow samples **(Figure 5K)**. Strikingly, whereas 100% of hIgG1-treated controls yielded replication-competent virus in TZM-bl assays, 40% (4/10) of bone marrow samples from 15×sCD4-Fc-treated animals showed complete viral suppression below detection limits **(Figure 5L)**. Critically, hyperIL-15-Fc showed no significant reservoir reduction in any assay **(Figure 5J-L)**, mirroring the well-documented limitations of N-803 which, despite demonstrating latency reversal in vitro, has shown minimal reservoir reduction in both preclinical models and clinical trials.^31–33,35,57^ These results highlighting 15×sCD4-Fc’s unique capacity for reservoir clearance.

Together, these findings establish 15×sCD4-Fc as a uniquely effective therapeutic agent that simultaneously achieves targeted reservoir reactivation, enhanced HIV-1-specific immunity and substantial reservoir reduction, addressing critical limitations of current latency reversal and immunotherapeutic approaches.

## DISCUSSION

The quest for HIV cure necessitates strategies that seamlessly integrate latency reversal with immune-mediated clearance,^58,59^ a goal hampered by the challenge of spatiotemporal discordance between LRAs-induced reservoir exposure and immune recognition. Our engineered multifunctional fusion protein 15×sCD4-Fc overcomes this fundamental limitation through temporally and spatially coupled mechanisms, unifying three synergistic functions: (i) hyperIL-15-driven transcriptional reactivation of latent reservoirs, (ii) sCD4-mediated precision targeting of Env^+^ cells, and (iii) Fc-dependent effector cell engagement amplified by hyperIL-15’s immunostimulatory properties. Crucially, 15×sCD4-Fc achieves what standalone therapies cannot: concurrent latency reversal and immune-mediated clearance, thereby resolving the long-standing “activation-clearance disconnect” that has undermined previous “shock-and-kill” approaches. This tripartite synergy, demonstrated by an 86.2% and 93.8% reduction in replication-competent reservoir size in treated humanized mice and in PBMCs from PLWH, respectively, along with complete elimination of replication-competent reservoirs in 36% of PLWH-derived samples by a single molecular agent, offers a clinically actionable path toward HIV-1 remission.

The “shock-and-kill” paradigm represents the most widely investigated strategy for HIV-1 reservoir eradication, with LRAs constituting its essential component.^60,61^ While most LRAs can reactivate latent HIV-1 in primary human CD4^+^ T cells, the reactivated cells often evade CD8^+^ T cell recognition,^25^ severely compromising their therapeutic efficacy. In contrast, IL-15 and its superagonists (e.g., N-803) not only reactivate latent virus but also sensitize reactivated reservoir cells to CD8^+^ T cell recognition in vitro while enhancing immune effector functions,^25^ making them promising candidates currently under clinical investigation.^35,62^ However, their clinical translation has been hindered by two fundamental limitations: inadequate reservoir reactivation outcomes in vivo, as demonstrated in SIV/HIV-1 animal models and human clinical trials,^31–33,35,57^ and failure to achieve significant reservoir reduction in SIV/HIV-1 infected animals^31,32^ and in people living with HIV-1^35^. Notably, N-803 did not augment the SIV-DNA reduction in lymphoid tissues of infected macaques achieved by LRA (AZD5582) plus SIV Env-specific antibodies^63^, suggesting that simply combination N-803 with LRA failed to achieve a syngenetic effect.

Our engineered 15×sCD4-Fc overcomes these barriers through synergistic integration of three functional modules. The coordinated IL-15R signaling and sCD4-mediated Env engagement establish a novel “tag-and-activate” mechanism, wherein sCD4 binding spatially concentrates the molecule on reactivated reservoirs, prolonging IL-15 signaling and amplifying viral production (8.8-fold enhancement versus hyperIL-15 alone), suggesting 15×sCD4-Fc may address a key therapeutic gap. Moreover, unlike N-803’s isolated immune-stimulation property, 15×sCD4-Fc directly couples latency reversal with sCD4-Env-targeted immune attack through Fc-dependent NK engagement, as demonstrated by its unique capacity to deplete lymphoid reservoirs in humanized models and in PBMCs from PLWH. Critically, our design differs from current bispecific antibody approaches under investigation,^64^ including bispecific CD16 engagers (targeting HIV-1 Env and human CD16)^64,65^ or bispecific CD3 engagers (e.g., Env×CD3 DARTs).^66–69^ Unlike these bispecific formats that require co-administration with potent LRAs (e.g., PMA/ionomycin or protein kinase C agonists) to achieve reservoir cell killing in vitro, 15×sCD4-Fc intrinsically integrates latency reversal and targeted immune clearance within a single molecule. Moreover, the simultaneous activation and engagement of both NK cells and CD8^+^ T cells by 15×sCD4-Fc minimizes the likelihood of immune escape during HIV treatment, thereby optimizing therapeutic efficacy.

This heterodimer design of 15×sCD4-Fc resolves a paradox in CD4-based biologics: whereas conventional homodimeric sCD4-Fc causes systemic MHC-II^+^ cell depletion via Fc-driven ADCC, our heterodimeric architecture exploits monomeric sCD4’s lack of binding to MHC-II^70^ to achieve HIV-specific targeting without off-target toxicity. Our experimental data clearly demonstrate that monomeric sCD4 within 15×sCD4-Fc enables selective Env engagement while avoiding promiscuous MHC-II interactions, a design principle that could inform future HIV-1 therapeutics. Although the neutralization potency of sCD4 for Env is lower than that of bNAbs, its broader binding capacity across diverse strains^71^, attributable to CD4’s natural role as the HIV-1 receptor, suggests that 15×sCD4-Fc may target the majority of reactivated reservoir cells in PLWH. In addition, engagement of the sCD4 molecule with HIV-1 Env on the surface of infected cells triggers Env to adopt an “open” conformation, thereby promoting endogenous antibody binding and subsequent ADCC.^39,72^ The resultant molecule 15×sCD4-Fc thus not only emulate the precision of bNAbs, with sCD4 conferring Env-binding specificity and Fc domain facilitating effector recruitment, but also harness endogenous antiviral antibody responses within the host. By combining the latency reversal activity of hyperIL-15, the 15×sCD4-Fc thereby creates a self-contained therapeutic circuit that overcomes the temporal and spatial discordance of conventional “shock-and-kill” approaches. In summary, our proof-of-concept study establishes tripartite integration of viral reactivation, precision targeting, and immune clearance as a transformative therapeutic paradigm against persistent HIV-1 reservoirs. To our knowledge, this constitutes a previously unreported engineered platform unifying three fundamental pillars of HIV cure strategies within a single molecular architecture, yielding synergistic outcomes. The inherent modularity of this system further enables tailored combinations with complementary latency-reversing agents or immune modulators.

## Limitations of the study

Our study establishes 15×sCD4-Fc as a multifunctional therapeutic agent coordinating HIV-1 reservoir reactivation and Env-directed immune clearance. However, there are several limitations that should be noted. First, the human immune system reconstituted in hu-mice is not fully functional compared to immunocompetent hosts.^73^ Although 15×sCD4-Fc enhances anti-HIV-1 immunity in hu-mice, the response may remain insufficient to fully control or eliminate the viral reservoir. We hypothesize that in immunocompetent hosts, intact lymphoid niches supporting robust innate and adaptive immunity would further improve reservoir clearance efficacy. Second, to avoid potential graft-versus-host disease (GVHD) in hu-mice beyond 6-9 months post-reconstitution,^73,74^ we terminated the experiment before 6 months, precluding long-term HIV-1 rebound assessment after cART cessation. Nevertheless, a recent macaque study demonstrated that dual immunotherapy (N-803 plus bNAbs) in SHIV-infected, ART-suppressed macaques achieved sustained viral control despite limited SHIV reservoir reduction.^57^ This suggests that 15×sCD4-Fc, which combines synergistic reservoir reactivation, targeted immune engagement and effector-mediated clearance function in one molecule, may outperform single molecule or simple combinatorial approaches, as evidenced by its significant reduction of replication-competent HIV-1 reservoirs both in cells from PLWH and in HIV-1 infected humanized mice. Third, although the hyper-IL-15 analog N803 has demonstrated safety and tolerability in clinical trials,^35,75^ the neoantigenic potential of our chimeric protein remains a concern. Current models mask this risk due to the compromised humoral immunity of humanized mice; thus, Phase I trials should incorporate immunopeptidomics screening to preempt antidrug antibody development.

## RESOURCE AVAILABILITY

### Lead contact

Further information and requests should be directed to the lead contact, Dr. Liang Cheng.

### Materials and data availability

All unique/stable reagents generated in this study are available from the lead contact with a completed materials transfer agreement. All data are available in the main text or the supplementary materials.

## ACKNOWLEDGMENTS

We acknowledge the support of the Biosafety Level 3 Laboratories of Infectious Disease Center, Guangzhou Eighth People’s Hospital. We gratefully acknowledge the BioBank of Guangzhou Eighth People’s Hospital for providing biosamples and services. We are also thankful to Dr. Liguo Zhang, Dr. Shengdian Wang and Dr. Ying Li from the Institute of Biophysics, Chinese Academy of Sciences and Dr. Lishan Su from University of Maryland for their valuable suggestions and discussions. Special thanks are extended to Ms. Lian Peng for her technical assistance. This work was supported by grants and fellowships from National Key R&D Program (2023YFC2306600 & 2024YFA1306500 to L.C.), National Natural Science Foundation of China (82572560 to L.C.), Fundamental Research Funds for the Central Universities (2042022dx0003 to L.W. and L.C.), Natural Science Foundation of Wuhan (2024040701010031 to L.C.), Wuhan Special Project for Knowledge Innovation (2023020201010070 to L.C.), Natural Science Foundation of Guangdong Province (2022A1515010589 to H.Y.) and Key-Area Research and Development Program of Guangdong Province (2022B1111020002 to H.Y.).

## AUTHOR CONTRIBUTIONS

Conceptualization: L.C., F.L.; Methodology: L.C., H.Y., F.L., L.W.; Investigation: F.L., Y.C., N.L., X.W., W.H., X.X., D.J., Y.X., J.W., H.W., Z.Y., P.Q., S.W., Z.L., T.C., S.S., Y.X., X.T., L.L., L.W.; Visualization: L.C., H.Y., L.F., L.W.; Funding acquisition: L.C., H.Y., L.W.; Project administration: L.C., H.Y.; Supervision: L.C., H.Y., L.W.; Writing – original draft: L.C., F.L.; Writing – review & editing: L.C., F.L., L.W., H.Y.

## SUPPLEMENTAL INFORMATION

Document S1. Figures S1-S6 and Tables S1-S3

## STAR METHODS

### KEY RESOURCES TABLE

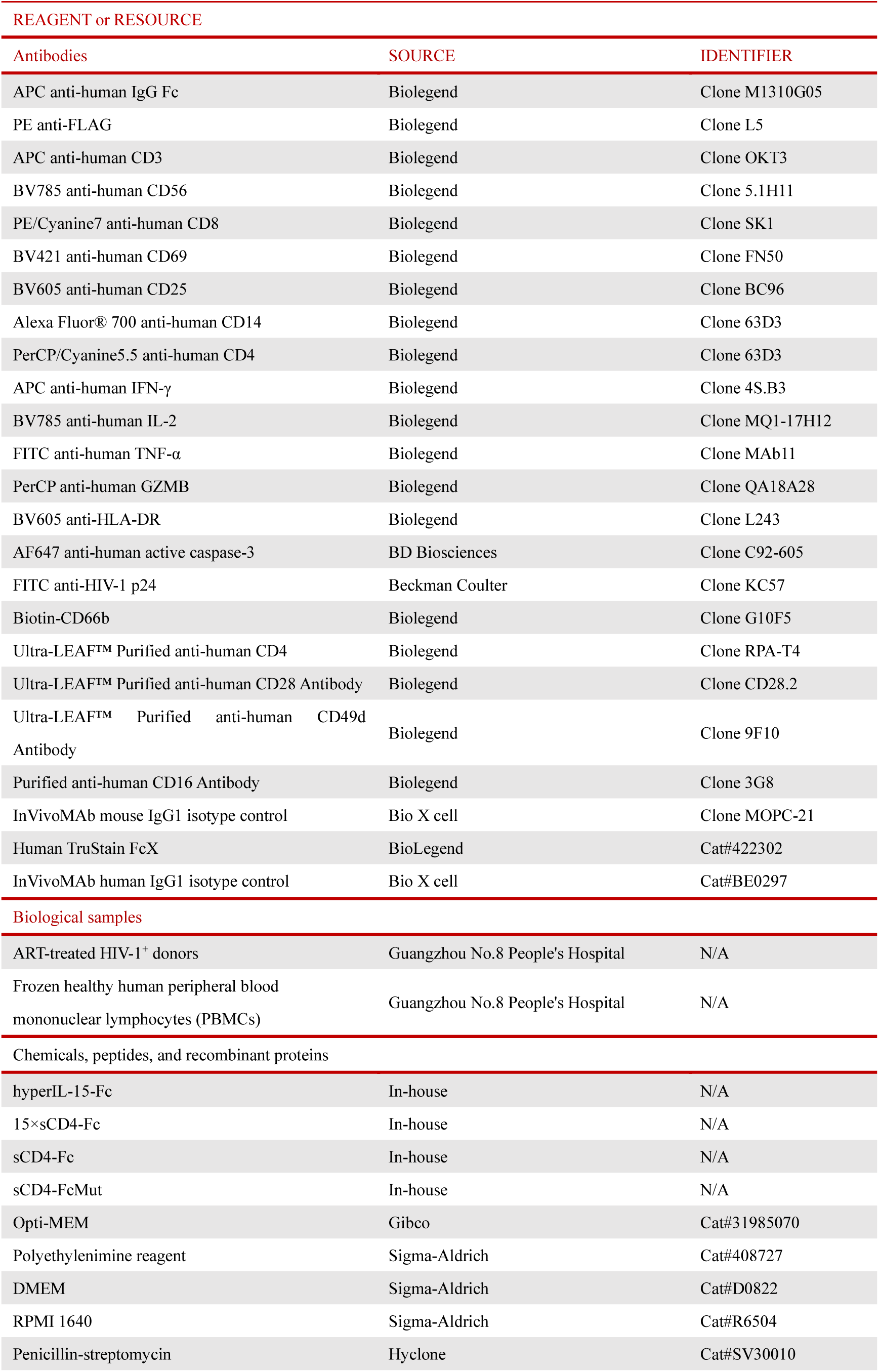
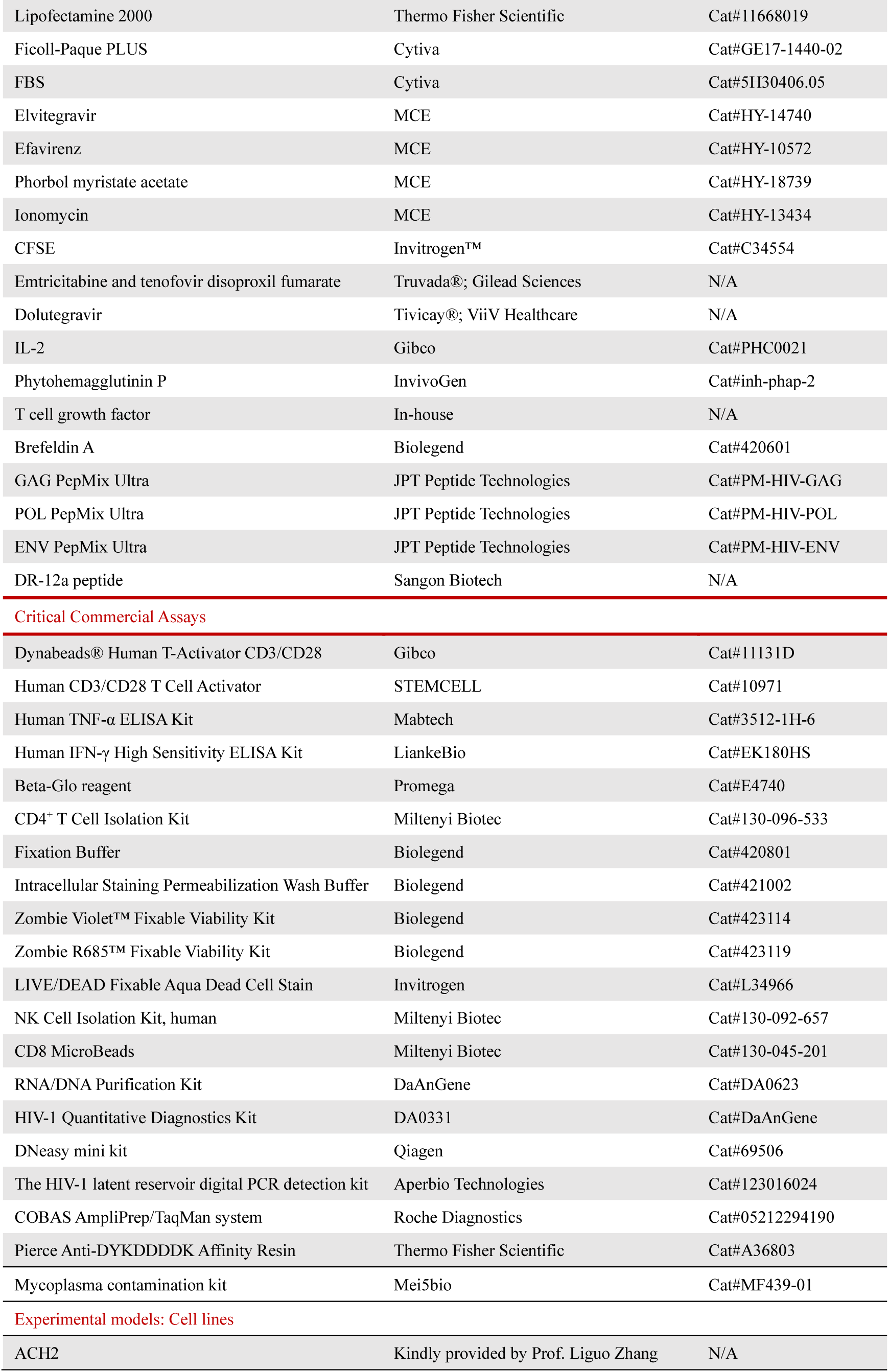
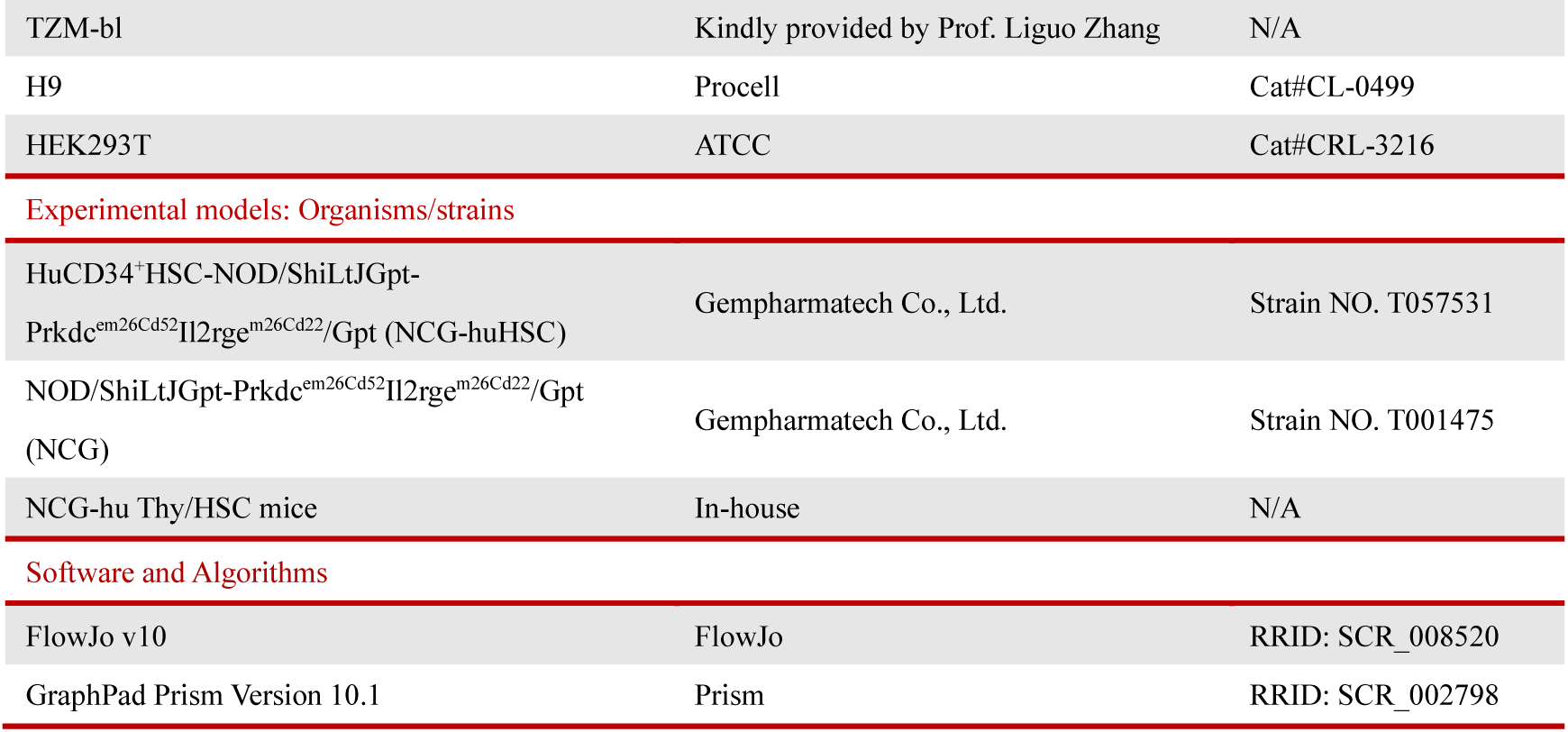

| REAGENT or RESOURCE |  |  |
| --- | --- | --- |
| Antibodies | SOURCE | IDENTIFIER |
| APC anti-human IgG Fc | Biolegend | Clone M1310G05 |
| PE anti-FLAG | Biolegend | Clone L5 |
| APC anti-human CD3 | Biolegend | Clone OKT3 |
| BV785 anti-human CD56 | Biolegend | Clone 5.1H11 |
| PE/Cyanine7 anti-human CD8 | Biolegend | Clone SK1 |
| BV421 anti-human CD69 | Biolegend | Clone FN50 |
| BV605 anti-human CD25 | Biolegend | Clone BC96 |
| Alexa Fluor® 700 anti-human CD14 | Biolegend | Clone 63D3 |
| PerCP/Cyanine5.5 anti-human CD4 | Biolegend | Clone 63D3 |
| APC anti-human IFN-γ | Biolegend | Clone 4S.B3 |
| BV785 anti-human IL-2 | Biolegend | Clone MQ1-17H12 |
| FITC anti-human TNF-α | Biolegend | Clone MAb11 |
| PerCP anti-human GZMB | Biolegend | Clone QA18A28 |
| BV605 anti-HLA-DR | Biolegend | Clone L243 |
| AF647 anti-human active caspase-3 | BD Biosciences | Clone C92-605 |
| FITC anti-HIV-1 p24 | Beckman Coulter | Clone KC57 |
| Biotin-CD66b | Biolegend | Clone G10F5 |
| Ultra-LEAF™ Purified anti-human CD4 | Biolegend | Clone RPA-T4 |
| Ultra-LEAF™ Purified anti-human CD28 Antibody | Biolegend | Clone CD28.2 |
| Ultra-LEAF™ Purified anti-human CD49d Antibody | Biolegend | Clone 9F10 |
| Purified anti-human CD16 Antibody | Biolegend | Clone 3G8 |
| InVivoMAb mouse IgG1 isotype control | Bio X cell | Clone MOPC-21 |
| Human TruStain FcX | BioLegend | Cat#422302 |
| InVivoMAb human IgG1 isotype control | Bio X cell | Cat#BE0297 |
| Biological samples |  |  |
| ART-treated HIV-1<sup>+</sup> donors | Guangzhou No.8 People's Hospital | N/A |
| Frozen healthy human peripheral blood mononuclear lymphocytes (PBMCs) | Guangzhou No.8 People's Hospital | N/A |
| Chemicals, peptides, and recombinant proteins |  |  |
| hyperIL-15-Fc | In-house | N/A |
| 15×sCD4-Fc | In-house | N/A |
| sCD4-Fc | In-house | N/A |
| sCD4-FcMut | In-house | N/A |
| Opti-MEM | Gibco | Cat#31985070 |
| Polyethylenimine reagent | Sigma-Aldrich | Cat#408727 |
| DMEM | Sigma-Aldrich | Cat#D0822 |
| RPMI 1640 | Sigma-Aldrich | Cat#R6504 |
| Penicillin-streptomycin | Hyclone | Cat#SV30010 |

| REAGENT or RESOURCE | SOURCE | IDENTIFIER |
| --- | --- | --- |
| Lipofectamine 2000 | Thermo Fisher Scientific | Cat#11668019 |
| Ficoll-Paque PLUS | Cytiva | Cat#GE17-1440-02 |
| FBS | Cytiva | Cat#5H30406.05 |
| Elvitegravir | MCE | Cat#HY-14740 |
| Efavirenz | MCE | Cat#HY-10572 |
| Phorbol myristate acetate | MCE | Cat#HY-18739 |
| Ionomycin | MCE | Cat#HY-13434 |
| CFSE | Invitrogen™ | Cat#C34554 |
| Emtricitabine and tenofovir disoproxil fumarate | Truvada®; Gilead Sciences | N/A |
| Dolutegravir | Tivicay®; ViiV Healthcare | N/A |
| IL-2 | Gibco | Cat#PHC0021 |
| Phytohemagglutinin P | InvivoGen | Cat#inh-phap-2 |
| T cell growth factor | In-house | N/A |
| Brefeldin A | Biolegend | Cat#420601 |
| GAG PepMix Ultra | JPT Peptide Technologies | Cat#PM-HIV-GAG |
| POL PepMix Ultra | JPT Peptide Technologies | Cat#PM-HIV-POL |
| ENV PepMix Ultra | JPT Peptide Technologies | Cat#PM-HIV-ENV |
| DR-12a peptide | Sangon Biotech | N/A |
| *Critical Commercial Assays* |  |  |
| Dynabeads® Human T-Activator CD3/CD28 | Gibco | Cat#11131D |
| Human CD3/CD28 T Cell Activator | STEMCELL | Cat#10971 |
| Human TNF-α ELISA Kit | Mabtech | Cat#3512-1H-6 |
| Human IFN-γ High Sensitivity ELISA Kit | LiankeBio | Cat#EK180HS |
| Beta-Glo reagent | Promega | Cat#E4740 |
| CD4<sup>+</sup> T Cell Isolation Kit | Miltenyi Biotec | Cat#130-096-533 |
| Fixation Buffer | Biolegend | Cat#420801 |
| Intracellular Staining Permeabilization Wash Buffer | Biolegend | Cat#421002 |
| Zombie Violet™ Fixable Viability Kit | Biolegend | Cat#423114 |
| Zombie R685™ Fixable Viability Kit | Biolegend | Cat#423119 |
| LIVE/DEAD Fixable Aqua Dead Cell Stain | Invitrogen | Cat#L34966 |
| NK Cell Isolation Kit, human | Miltenyi Biotec | Cat#130-092-657 |
| CD8 MicroBeads | Miltenyi Biotec | Cat#130-045-201 |
| RNA/DNA Purification Kit | DaAnGene | Cat#DA0623 |
| HIV-1 Quantitative Diagnostics Kit | DA0331 | Cat#DaAnGene |
| DNeasy mini kit | Qiagen | Cat#69506 |
| The HIV-1 latent reservoir digital PCR detection kit | Aperbio Technologies | Cat#123016024 |
| COBAS AmpliPrep/TaqMan system | Roche Diagnostics | Cat#05212294190 |
| Pierce Anti-DYKDDDDK Affinity Resin | Thermo Fisher Scientific | Cat#A36803 |
| Mycoplasma contamination kit | Mei5bio | Cat#MF439-01 |
| *Experimental models: Cell lines* |  |  |
| ACH2 | Kindly provided by Prof. Liguozhang | N/A |

| *Continued* |  |  |
| --- | --- | --- |
| REAGENT or RESOURCE | SOURCE | IDENTIFIER |
| TZM-bl | Kindly provided by Prof. Liguozhang | N/A |
| H9 | Procell | Cat#CL-0499 |
| HEK293T | ATCC | Cat#CRL-3216 |
| **Experimental models: Organisms/strains** |  |  |
| HuCD34<sup>+</sup>HSC-NOD/ShiLtJGpt-Prkdc<sup>em26Cd52</sup>Il2rge<sup>m26Cd22</sup>/Gpt (NCG-huHSC) | Gempharmatech Co., Ltd. | Strain NO. T057531 |
| NOD/ShiLtJGpt-Prkdc<sup>em26Cd52</sup>Il2rge<sup>m26Cd22</sup>/Gpt (NCG) | Gempharmatech Co., Ltd. | Strain NO. T001475 |
| NCG-hu Thy/HSC mice | In-house | N/A |
| **Software and Algorithms** |  |  |
| FlowJo v10 | FlowJo | RRID: SCR_008520 |
| GraphPad Prism Version 10.1 | Prism | RRID: SCR_002798 |

### EXPERIMENTAL AND STUDY PARTICIPANT DETAILS

#### Human participants

Primary human PBMCs were obtained from ART-treated PLWH donors at Guangzhou No.8 People’s Hospital with ethical approval by the Medical Ethics Committee (202033166).

#### Construction of humanized mice

NCG-hu Thy/HSC mice were generated using our previously established protocol.^76^ In brief, human fetal liver and thymus tissues (gestational age 16-20 weeks) were obtained from elective or medically indicated pregnancy terminations through Guangzhou Women and Children’s Medical Center, with written informed consent obtained from all maternal donors. Six-week-old NCG mice (NOD/ShiLtJGpt-Prkdc^em26Cd52^Il2rge^m26Cd22^/Gpt) underwent sublethal irradiation followed by anesthesia. Human fetal thymus fragments were surgically implanted under the renal capsule, while CD34^+^ hematopoietic progenitor cells isolated from matched fetal liver tissues were immediately administered intravenously (1 × 10^5^ cells/mouse). Human immune cell engraftment was assessed by flow cytometry 12 weeks post-transplantation. All procedures were approved by the Guangzhou Women and Children’s Medical Center Institutional Review Board (determined non-human subjects research, approval #[2024]NO.059A01). Commercially available humanized NCG mice reconstituted with intravenous-administered human cord blood-derived CD34^+^ hematopoietic cells (NCG-hu HSC mice) were obtained from Gempharmatech Co., Ltd. All animal procedures were conducted in compliance with approved guidelines by the Guangzhou No.8 People’s Hospital Institutional Animal Care and Use Committee (gz8h-YHS-2023-03).

## METHOD DETAILS

### Construction and expression of fusion proteins

The expression cassettes for hyperIL-15-Fc, sCD4-Fc, and 15×sCD4-Fc were shown in Figure S1. The hyperIL-15 component was generated by fusing IL-15 to the IL-15Rα sushi domain via a SGGSGGGGSGGGSGGGGSLQ linker. hyperIL-15-Fc was constructed by linking this hyperIL-15 module to the hIgG1-Fc domain through an additional 3×GGGGS linker and cloned into the pTT3 vector. sCD4-Fc was produced by connecting the D1D2 domain of human CD4 to hIgG1-Fc using a 3×GGGGS linker and cloned into the pTT3 vector. sCD4-FcMut was constructed by engineered with ‘P329G/L234A/L235A’ mutants into CH2 domains of hIgG1-Fc to reduce Fc receptor (FcR) binding.^48^ For 15×sCD4-Fc construction, hyperIL-15 and sCD4 were independently cloned into pEE6.4-Fc6 and pEE6.4-Fc9 plasmids (Lonza), respectively, each incorporating a 3×GGGGS linker between the functional domain (hyperIL-15 or sCD4) and the corresponding Fc variant (hIgG1-Fc6 or hIgG1-Fc9). Heterodimerization of hyperIL-15 and sCD4 and Fc was achieved by the knobs-into-holes approach.^77,78^ The constructions hyperIL-15-Fc, sCD4-Fc, and sCD4-Fc9 included a C-terminal 3×FLAG tag.

Protein expression was performed by transient transfection of plasmids into HEK 293T cells, where hyperIL-15-Fc, sCD4-Fc or sCD4-FcMut were transfected individually, while 15×sCD4-Fc was produced by co-transfecting hyperIL-15-Fc6 and sCD4-Fc9-FLAG plasmids at a 1:1 ratio with heterodimer formation enabled through knob-into-hole interactions between hIgG1-Fc6 and hIgG1-Fc9 domains. Briefly, plasmid(s) was diluted into Opti-MEM, mixed with polyethylenimine reagent (Sigma, #408727) for 15 min at room temperature, and added into HEK 293T cells. 48-72 hours post transfection, the supernatant from transfected cells was collected and filtered through 0.45 µm filter unit (Millipore, #HPWP01300). The protein in the supernatant was purified using Pierce Anti-DYKDDDDK Affinity Resin (Thermo Fisher Scientific, #A36803) and the bound protein was eluted using the glycine buffer (100 mM, PH=2.8). The eluted product was dialyzed against PBS (pH7.4) overnight with 4 changes of buffer. The purified protein was analyzed on non-reduced and reduced SDS-PAGE.

### Flow cytometric analysis

For cell surface marker analysis, single-cell suspensions were washed twice with PBS containing 2% FBS. Cells were then incubated with Human TruStain FcX (BioLegend, #422302) at 4 °C for 10 minutes to block Fc receptors, followed by staining with Viability Kit for 10 minutes to assess cell viability. Surface marker staining was performed by incubating cells with fluorochrome-conjugated antibodies for 30 minutes at 4 °C. For intracellular staining, cells were first stained with surface markers and then fixed and permeabilized with Fixation Buffer (Biolegend, #420801) and Intracellular Staining Permeabilization Wash Buffer (Biolegend, #421002) followed by intracellular staining. After staining, cells were washed and resuspended in 1% paraformaldehyde (PFA) for acquisition on a BD LSRFortessa X-20 flow cytometer (BD Biosciences). Data were analyzed using FlowJo software (v10.8.1, BD Biosciences).

### Validation of the bioactivity of fusion proteins

The immunostimulatory activity of the fusion proteins was assessed by culturing human PBMCs with 10 nM of hIgG1 control, hyperIL-15-Fc, or 15×sCD4-Fc for 48 hours, followed by flow cytometric analysis of CD69 expression on CD4^+^ and CD8^+^ T cells, and NK cells.

Neutralization activity was evaluated using TZM-bl assays, where HIV-1_JRCSF_ (140 PFU) was pre-incubated with serially diluted (100∼0.032 μg/mL) hyperIL-15-Fc, sCD4-Fc, or 15×sCD4-Fc in complete DMEM for 1 hour at 37 °C before addition to 1×10^4^ TZM-bl cells and further cultured for 44 hours. Luciferase activity in TZM-bl cells was measured by M5 Luciferase Reporter Assay Kit (Mei5bio, # MF489-01). Briefly, after removal of supernatant, cells were lysed with 50 μL lysis buffer and centrifuged at 12,000 × g for 5 minutes to clear debris, with luciferase activity quantified using 100 μL luciferin buffer on a SpectraMax i3x system (Molecular Devicesmd). HIV-1 virus stocks used in the study were generated by transfecting 293T cells with the JR-CSF infectious molecular clone (pYK-JRCSF) using Lipofectamine 2000 (Thermo Fisher Scientific, #11668019). Virus was titrated by infected TZM-bl cells by counted blue-stained cells after β-galactosidase activity staining.

Fc receptor binding activity was determined by incubating human PBMCs with hyperIL-15-Fc or 15×sCD4-Fc for 20 minutes at room temperature, with FcR blocking performed by pre-incubation with human serum for 10 minutes. After washing with 2% FPBS, cells were stained with anti-CD3, -CD56, -CD14, and -FLAG antibodies for 20 minutes and analyzed by flow cytometry.

### Reactivation of HIV-1 reservoir in primary CD4^+^ T cells from PLWH

cART-suppressed PLWH participants were selected based on criteria including continuous plasma viral load suppression (<20 copies/mL for >12 months), CD4^+^ T cell counts (>500 cells/μL), and absence of hepatitis B or C virus co-infection. Peripheral blood mononuclear cells (PBMCs) were isolated from freshly drawn blood samples using Ficoll-Paque PLUS (Cytiva) density gradient centrifugation at 900 × g for 30 minutes at room temperature with brake disabled. CD4^+^ T cells were negatively isolated using the CD4^+^ T Cell Isolation Kit, human (Miltenyi Biotec, #130-096-533) with supplemental biotinylated anti-human CD66b antibody (BioLegend, clone G10F5). Magnetic separation was performed using LS columns (Miltenyi Biotec) with a manual MACS separator (Miltenyi Biotec) according to the manufacturer’s protocol. The purity of CD4^+^ T cells was identified by flow cytometry.

The purified CD4^+^ T cells (purity >99% by flow cytometry) were immediately used for reservoir activation assays. For these assays, 3 × 10⁶ CD4^+^ T cells were plated per well in 12-well tissue culture plates (Corning) containing 1 mL complete RPMI 1640 medium and antiretroviral drugs elvitegravir (1 μM, MCE) and efavirenz (1 μM, MCE) to prevent new rounds of infection. For blocking HIV gp120 binding site in sCD4, 50 nM Ultra-LEAF™ Purified anti-human CD4 (BioLegend, clone RPA-T4) or isotype (Bio X cell, #BE0083) were incubated with 10 nM 15×sCD4-Fc 20 minutes in room temperature. Test proteins (hyperIL-15-Fc, 15×sCD4-Fc, 15×sCD4-Fc with anti-CD4 or hIgG1 isotype control) were added at 10 nM concentrations for 72 hours. To quantify HIV-1 RNA levels, 1 mL of culture supernatant was collected and analyzed using the robotic COBAS AmpliPrep/TaqMan system (Roche Diagnostics) with a validated detection limit of 20 copies/mL according to manufacturer’s protocols. Cell activation was stained with anti-CD69 and anti-CD25 by flow cytometric.

### Targeting assay of fusion proteins to activated ACH2 cells

Latently HIV-1-infected ACH2 cells were reactivated by 48-hour stimulation with 10 ng/mL phorbol myristate acetate (PMA), while non-activated controls were maintained in parallel. Following reactivation, both PMA-activated and non-activated ACH2 cells were incubated with 20 nM of hyperIL-15-Fc or 15×sCD4-Fc at room temperature for 20 minutes. After washing with 2% FPBS, cells were stained with anti-FLAG antibody and analyzed by flow cytometry to quantify fusion protein binding specificity. To block fusion protein targeting, 20 nM sCD4-Fc or 15×sCD4-Fc were incubated with 100 nM Ultra-LEAF™ Purified anti-human CD4 (BioLegend, clone RPA-T4) for 20 minutes at room temperature prior to 20-minute incubation with PMA-activated ACH2 cells.

### Binding of fusion proteins to HLA-DR^+^ cells

HLA-DR expression on H9 cells was confirmed by flow cytometry prior to binding assays. For functional testing, H9 cells were incubated with either 0.1 μg sCD4-Fc or 15×sCD4-Fc in 50 μl 2% FPBS (PBS containing 2%FBS) at 37 °C for 3 hours. Selected conditions included 50 μg/mL DR-12a peptide (EEYVRFDSDVGE, corresponding to amino acids 35-46 of the HLA-DR β chain; Sangon Biotech, ≥96% purity) as a competitive inhibitor. Following incubation, cells were washed with 1% FPBS and stained with anti-FLAG antibody for 30 minutes at 4°C. Protein binding to H9 cells was quantified by flow cytometry.

### ADCC Assays for evaluating fusion protein activity

The capacity of fusion proteins to mediate ADCC against HIV-1^+^ cells was assessed by co-culturing PMA-activated ACH2 cells (targets) with primary human NK cells (effectors) in the presence of test proteins. NK cells were isolated from healthy donor PBMCs by using the NK Cell Isolation Kit, human (Miltenyi Biotec, #130-092-657) and served as effector cells. Target cells (PMA-activated ACH2) and control cells (non-activated ACH2) were differentially labeled with 5 μM or 0.5 μM CFSE, respectively. Effector and target cells were co-cultured at a 30:1 ratio in 96-well U-bottom plates with either 100 nM control hIgG1, hyperIL-15-Fc, or 15×sCD4-Fc and incubation for 15 minutes at room temperature followed by centrifugation at 300 × g for 1 minute. Plates were then incubated at 37 °C, 5% CO_2_ for 5 hours before Guava flow cytometry analysis (Cytek® Guava® Systems). The percentage of cytotoxicity was calculated with the following formula: [1-(Ratio (Target cell counts/ control cell counts) _+effector_ / Ratio (Target cell counts/ control cell counts) _-effector_)] × 100%. The cytotoxicity percentages observed in the hIgG1 control group served as background values and were systematically subtracted from all experimental groups during final data analysis.

For HLA-DR^+^ H9 cell ADCC assays, CFSE-labeled (1 μM) H9 cells were preincubated with 100 nM test proteins for 3 hours. Where indicated, sCD4-Fc was pre-blocked with 100 μg/mL DR-12a (20 minutes at RT) prior to adding NK cells (effector:target = 30:1) in 96-well U-bottom plates (Corning). After 15 minutes RT incubation followed by centrifugation at 300 × g for 1 minute, cells were cultured at 37 °C, 5% CO_2_ for 6 hours before Guava flow cytometry analysis (Cytek® Guava® Systems). The percentage of cytotoxicity was calculated with the following formula: [1-Ratio (H9 cell counts_+effector_ / H9 cell counts_–effector_)] × 100%. The cytotoxicity percentages observed in the hIgG1 control group served as background values and were systematically subtracted from all experimental groups during final data analysis.

### CTL assay for fusion protein activity evaluation

HIV-1-infected participants were selected based on criteria including continuous plasma viral load suppression (<20 copies/mL for >12 months), CD4^+^ T cell counts (>500 cells/μL), and absence of hepatitis B or C virus co-infection. CD4^+^ T cells were isolated using the CD4^+^ T Cell Isolation Kit, human (Miltenyi Biotec, #130-096-533) and superinfected with HIV-1_JR-CSF_ (MOI = 0.2) for 60 hours. Autologous CD8^+^ T cells were selected using CD8 MicroBeads (Miltenyi Biotec, #130-045-201) and pre-treated with 10 nM hIgG1 or 15×sCD4-Fc for 60 hours. The cells were then washed and cocultured at a 20:1 effector-to-target ratio (CD8^+^:CD4^+^) in 96-well U-bottom plates for 6 hours. Cell numbers were quantified by flow cytometry (Cytek® Guava® Systems) followed by surface marker staining (CD3/CD4/CD8) and intracellular detection (p24/active caspase-3).

### Ex vivo latency eradication assay

HIV-1-infected participants were selected based on criteria including continuous plasma viral load suppression (<20 copies/mL for >12 months), CD4^+^ T cell counts (>500 cells/μL), and absence of hepatitis B or C virus co-infection. PBMCs from ART-suppressed PLWH were cultured with 10 nM 15×sCD4-Fc or hIgG1 control in media containing efavirenz (300 nM) and IL-2 (10 U/mL) for 7 days. On day 3, CD8^+^ T-cell and NK cell functionality were assessed. For HIV-1 specific CD8^+^ T cell responses, cells were stimulated with pooled HIV-1 peptides (GAG/POL/ENV PepMix Ultra; 2 μg/mL per peptide; JPT Peptide Technologies) for 3 hours without brefeldin A, followed by 5 hours incubation with brefeldin A prior to surface and intracellular cytokine staining using established protocols. NK cell function was evaluated by stimulating cells with plate-bound anti-CD16 antibody (clone 3G8; BioLegend, #302001) for 1 hour without brefeldin A, followed by 4-hour brefeldin A treatment before flow cytometry.

On day 7, CD4^+^ T cells were isolated using CD4^+^ T Cell Isolation Kit, human (Miltenyi Biotec, #130-096-533) with supplemental biotinylated anti-human CD66b antibody (BioLegend, clone G10F5), followed by quantification of HIV *gag* DNA or replication-competent HIV-1 using TZM-bl reporter assays.

### HIV-1 infection of humanized mice

Humanized mice demonstrating stable human leukocyte reconstitution (>20% hCD45^+^ cells in peripheral blood) were anesthetized and infected via retro-orbital injection with 3,000 infectious units (IU) of HIV-1_JR-CSF_ per mouse in 30 μL PBS.

### Combination antiretroviral therapy (cART)

Antiretroviral drug-formulated food was prepared following established methods^50^ with modification by substituting raltegravir with dolutegravir. Briefly, commercial tablets of emtricitabine and tenofovir disoproxil fumarate (Truvada®; Gilead Sciences) and dolutegravir (Tivicay®; ViiV Healthcare) were pulverized into homogeneous powder and uniformly mixed with Irradiated Immunodeficient Lab Rat and Mouse Diet (XTI01MY-002). The final drug concentrations in formulated chow were: 260 mg/kg dolutegravir, 1,560 mg/kg tenofovir disoproxil, and 1,040 mg/kg emtricitabine. The estimated daily drug doses were 42 mg/kg dolutegravir, 250 mg/kg tenofovir disoproxil, and 166 mg/kg emtricitabine.

### In vivo fusion proteins treatments

In vivo fusion protein treatments were administered to HIV-1-infected, cART-treated humanized mice via intraperitoneal (i.p.) injection beginning at 7 weeks postinfection. Treatment groups received either sCD4-Fc, sCD4-FcMut, hyperIL-15-Fc, or 15×sCD4-Fc at doses of 20 μg/mouse twice weekly for 2-3 weeks (specific durations varied by experimental design as indicated in figure legends). All control groups received equivalent doses of human IgG1 isotype control (Bio X Cell, # BE0297).

### HIV-1 genomic RNA detection in plasma

HIV-1 genomic RNA was isolated from plasma using the RNA/DNA Purification Kit (DA0623, DaAnGene), a magnetic bead-based system for viral nucleic acid extraction. Quantification was performed by real-time PCR using the HIV-1 Quantitative Diagnostics Kit (DaAnGene, DA0331), with an assay sensitivity of 250 IU/mL (147 copies/mL) in plasma.

### Cell-associated HIV-1 DNA detection

To measure total cell-associated HIV-1 DNA, genome DNA was extracted from 1 × 10^6^ purified CD4^+^ T cells or 2 × 10^6^ human cells from spleen and bone marrow cells using the DNeasy mini kit (Qiagen). HIV-1 *gag* DNA was quantified by real-time PCR. DNA from serial dilutions of ACH2 cells, which contain 1 copy of HIV-1 genome in each cell, was used to generate a standard curve. Human CD4^+^ T cells numbers were determined by flow cytometry. For relative comparison, total human cell numbers were also determined by simultaneously amplifying human gamma globin DNA. HIV *gag* genome was detected with primers of 5′-GGTGCGAGAGCGTCAGTATTAAG-3′ and 5′-AGCTCCCTGCTTGCCCATA-3′. The probe (FAM-AAAATTCGGTTAAGGCCAGGGGGAAAGAA-QSY7) was ordered from Sangon Biotech and the reactions were set up following the manufacturer’s guidelines and were run on the CFX96 Real-Time PCR (BioRad). The sequences for the forward and reverse primers and probe for the detection of human gamma globin were 5’-CGCTTCTGGAACGTCTGAGATT-3’, 5’-CCTTGTCCTCCTCTGTGAAATGA-3’ and FAM-TCAATAAGCTCCTAGTCCAGAC-QSY7), respectively. Real-time PCR was performed as described above.

### Intact proviral DNA assay (IPDA)

The HIV-1 latent reservoir digital PCR detection kit (Aperbio Technologies, Suzhou, China) was used to quantify intact HIV-1 proviruses in spleen and bone marrow samples from HIV-1-infected humanized mice. This multiplex digital PCR assay simultaneously detects: (i) two distinct HIV-1 genomic regions (ψ and env) to differentiate intact versus defective proviruses, and (ii) two human RPP30 reference targets (5’RPP30 and 3’RPP30) for cellular DNA quantification and DNA shearing index (DSI) calculation, enabling precise normalization of proviral copy numbers per million cells. The intact HIV-1 copies were calculated according to the manufacturer’s instructions.

### TZM-bl assay (TZA) for quantification of inducible, replication-competent latent HIV-1

The replication-competent latent HIV-1 reservoir in purified CD4^+^ T cells from PLWH and in lymphoid organs of humanized mice was quantified using an optimized TZA adapted from established protocols.^49^ To quantify the replication-competent latent HIV-1 reservoir in CD4^+^ T cells from PLWH, the sorted CD4^+^ T cells were activated with Dynabeads® Human T-Activator CD3/CD28 (Gibco, #11131D) in the presence of efavirenz (300 nM) and IL-2 (50 U/mL) for 5 days. Following activation, cells were serially diluted four-fold (from 100,000 to 6,250 cells/well) and plated in octuplicate onto TZM-bl reporter cells (30,000-60,000 cells/well in RPMI-1640 medium supplemented with 10% FBS) in 96-well plates. After 48-hour coculture, CD4^+^ T cells were removed by washing, and TZM-bl monolayers were incubated with Beta-Glo reagent (Promega) for 1 hour before chemiluminescence measurement. CD4^+^ T cells isolated from PBMCs of heathy control donors were processed in parallel as negative controls.

For the replication-competent latent HIV-1 reservoir in lymphoid organs of humanized mice, bone marrow cells (human CD4^+^ T cells numbers were determined by flow cytometry) from HIV-1-infected hu-mice were activated with phytohemagglutinin (PHA, 2 μg/mL), IL-2 (100 U/mL), 1% T cell growth factor (prepared as previously described^79^) in the presence of efavirenz (EFV, 300 nM) for 48 hours. Following PHA removal, cells were maintained in fresh medium containing IL-2 (100 U/mL), 1% T cell growth factor, and EFV (300 nM) for an additional 48 hours. Activated BM cells were then serially diluted four-fold (100,000-6,250 cells/well) and plated in octuplicate onto TZM-bl reporter cells (30,000-60,000 cells/well in RPMI-1640 with 10% FBS) in 96-well plates. After 48-hour coculture, cells were removed by washing, and TZM-bl monolayers were incubated with Beta-Glo reagent (Promega) for 1 hour before chemiluminescence measurement. BM cells from uninfected hu-mice were processed in parallel as negative controls.

A well was considered positive if its luminescence exceeded the average value plus two times the standard deviation (mean + 2SD) of control wells containing either healthy donor cells or uninfected humanized mice samples. Infectious units per million (IUPM) cells were calculated using maximum likelihood estimation via the online IUPM statistical program (http://silicianolab.johnshopkins.edu/).

### T cell stimulation and intracellular cytokine staining

For nonspecific stimulation, splenocytes from hu-mice were cultured with phorbol 12-myristate 13-acetate (PMA; 50 ng/mL) and ionomycin (1 μM; MedChemExpress) in the presence of brefeldin A (1×; BioLegend, #420601) 6 hours. For T cell receptor (TCR) stimulation, splenocytes were treated with Human CD3/CD28 T Cell Activator (STEMCELL Technologies, #10971) for 3 hours without brefeldin A, followed by 5-hour stimulation with brefeldin A. Antigen-specific responses were assessed by incubating mixture of splenocytes and BM cells with pooled HIV-1 peptides (GAG/POL/ENV PepMix Ultra, 2 μg/mL per peptide; JPT Peptide Technologies) plus anti-CD28 (BioLegend, #302933) /CD49d (BioLegend, #304339) co-stimalation for 3 hours, followed by 5-hour brefeldin A treatment. Cells were then processed for surface and intracellular cytokine staining using established protocols. For supernatant cytokine analysis, mixture of splenocytes and BM cells were stimulated with HIV-1 peptide pools for 24 hours, after which supernatant was collected and analyzed for IFN-γ (Human IFN-γ High Sensitivity ELISA Kit, LiankeBio, #EK180HS) and TNF-α (Human TNF-α ELISA Kit, Mabtech, # 3512-1H-6) secretion.

### Statistics

All statistical analyses were performed using GraphPad Prism 10.1 (GraphPad Software). Data are presented as mean ± SEM or paired data plot. For comparisons between two groups, two-tailed Student’s t-tests were applied. Multiple group comparisons were analyzed by one-way ANOVA with Tukey’s post hoc test or non-parametric Kruskal-Wallis *H* test. Specific tests used for each dataset are detailed in corresponding figure legends. Significance thresholds were defined as \**P* < 0.05, \*\**P* < 0.01, \*\*\**P* < 0.001, and \*\*\*\**P* < 0.0001, with *P* value less than 0.05 was considered significant.

