## Supplementary material for "Synchronized Latency Reversal and Immune Clearance by a Multifunctional Fusion Protein Enables HIV-1 Reservoir Reduction": Figure S1-S6 and Table S1-S3

**The file includes:**

Figure S1 to S6

Tables S1 to S3

### Supplemental figures and legends

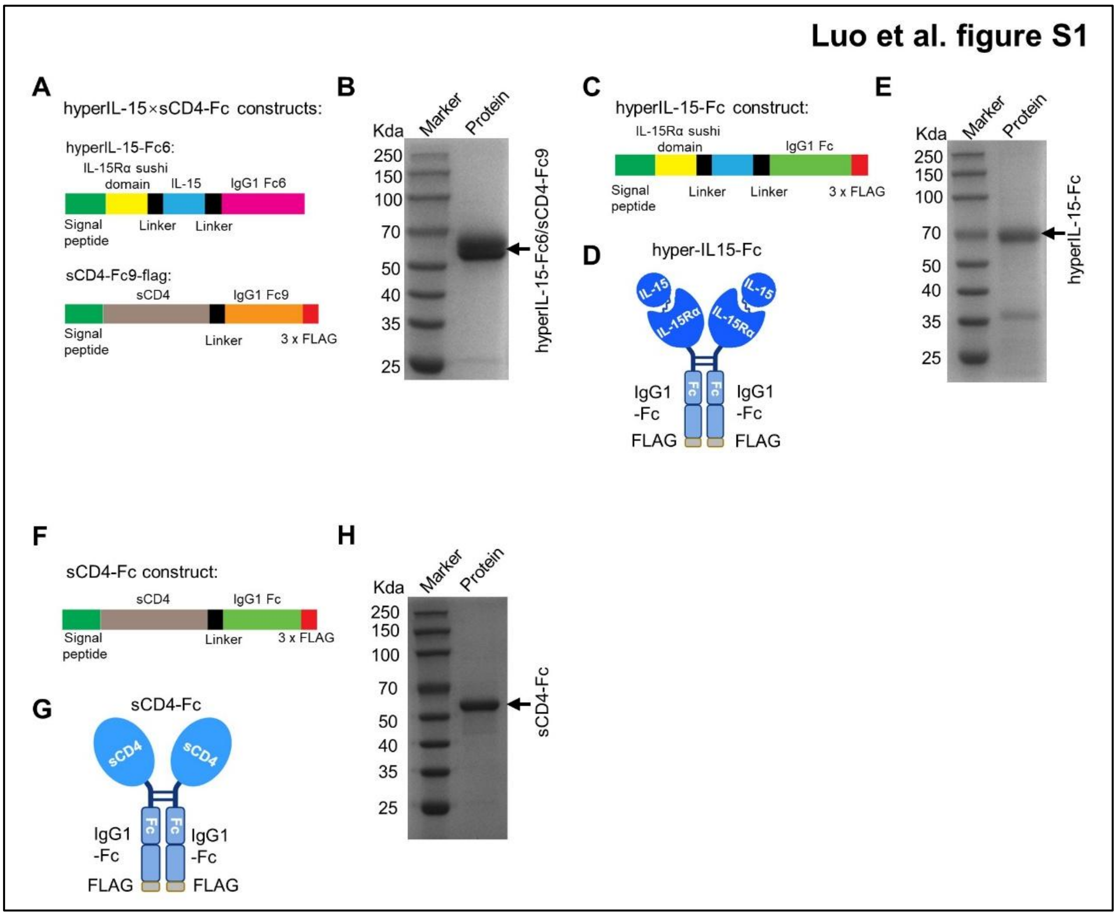


**Figure S1. Construction of 15×sCD4-Fc and control proteins**

**(A)** Schematic diagrams of the hyperIL-15-Fc6 and sCD4-Fc9 plasmids used for 15×sCD4-Fc heterodimer expression.

**(B)** SDS-PAGE analysis of purified 15×sCD4-Fc under reducing condition.

**(C)** Schematic diagram of the hyperIL-15-Fc plasmid construct.

**(D)** Schematic architecture of hyperIL-15-Fc.

**(E)** SDS-PAGE analysis of purified hyperIL-15-Fc under reducing conditions.

**(F)** Schematic diagram of the sCD4-Fc plasmid construct.

**(G)** Schematic architecture of sCD4-Fc.

**(H)** SDS-PAGE analysis of purified sCD4-Fc under reducing conditions.

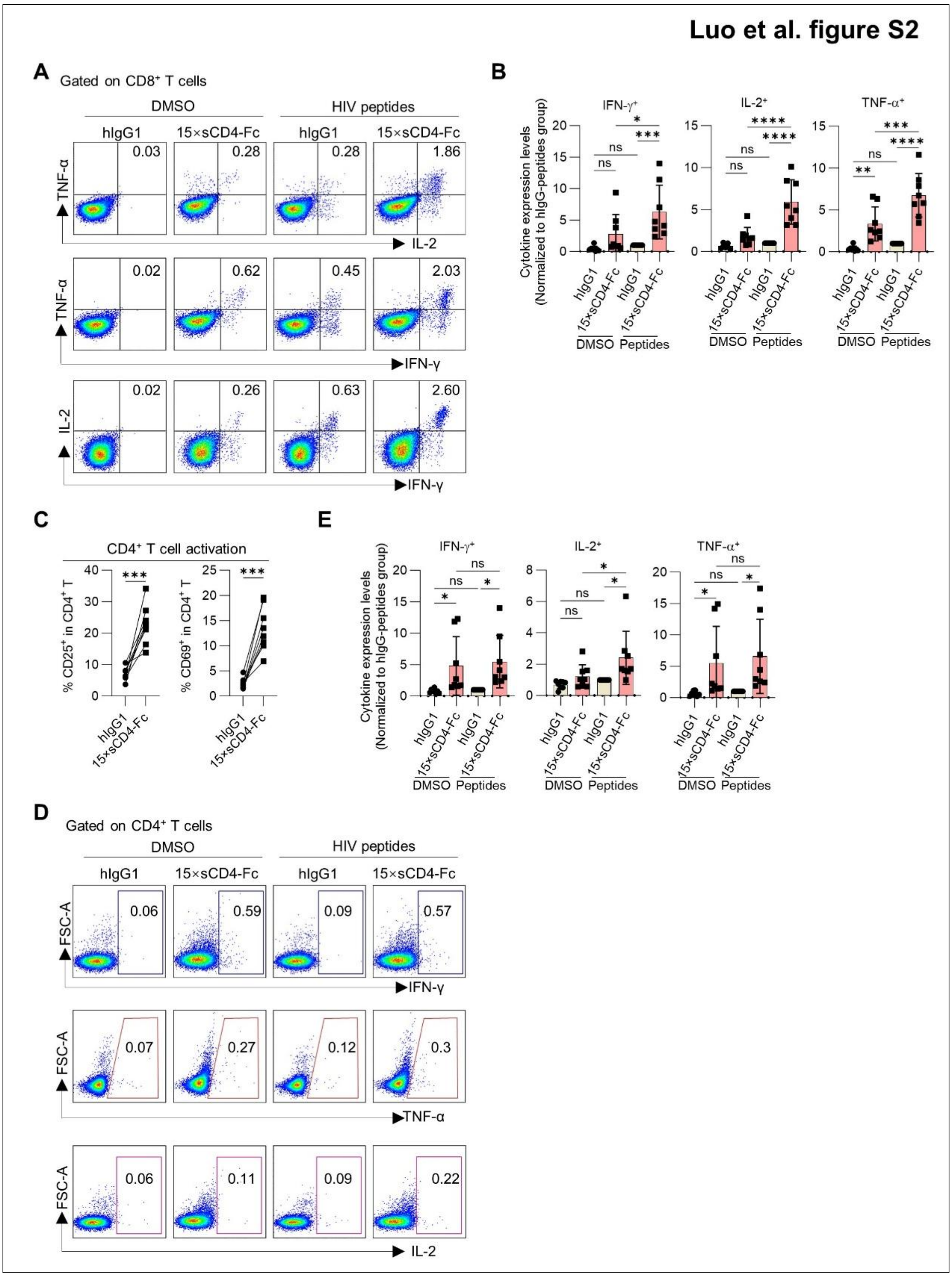


**Figure S2. The effects of 15×sCD4-Fc on HIV-1 specific CD8$^{+}$ and CD4$^{+}$ T cell response in PBMCs from ART-suppressed PLWH**

PBMCs from ART-suppressed PLWH (n=8 donors) were treated as in Figure. 1E.

**(A-B)** On day 3, PBMCs were stimulated with HIV-1 peptide pools for 8 hours (Brefeldin A added at 3 hours). Representation flow cytometry plots (A) and summary data (B) show expression of IFN-γ, IL-2 and TNF-α in CD8$^{+}$ T cells after HIV-1 peptide pools stimulation.

**(C)** Summary data show expression of CD25 and CD69 on CD4$^{+}$ T cells at day 3.

**(D-E)** On day 3, PBMCs were stimulated with HIV-1 peptide pools for 8 hours (Brefeldin A added at 3 hours). Representation flow cytometry plots (D) and summary data (E) show expression of IFN-γ, IL-2 and TNF-α in CD4$^{+}$ T cells after HIV-1 peptide pools stimulation.

Data in (B and E) are presented as mean ± SEM. Data in (C) are presented as paired data plot. Statistical significance was determined by one-way ANOVA followed by Tukey's post hoc multiple comparisons test (B and E) or paired Student's t-test (C). *$p < 0.05$, ***$p < 0.001$, ****$p < 0.0001$.

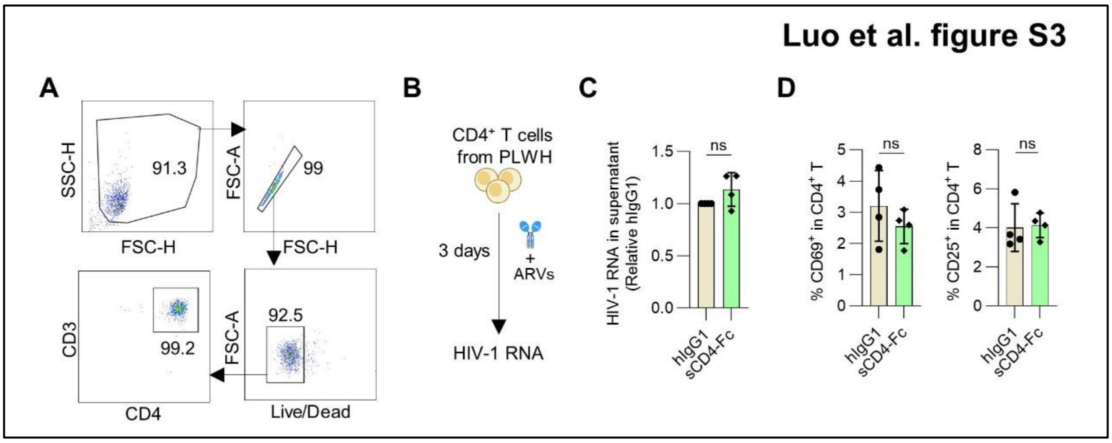


**Figure S3. sCD4-Fc lacks latency-reversing activity in primary CD4$^+$ T cells from PLWH**

**(A)** Representative flow cytometry gating strategy showing purity of CD4$^+$ T cells isolated from PBMCs of PLWH.

**(B)** Experimental schematic: primary CD4$^+$ cells isolated from cART-suppressed subjects were cultured for 72 hours with sCD4-Fc or hIgG1 (10 nM) in ARVs-containing media, followed by HIV-1 detection in supernatant and flow cytometric analysis of cells activation (n=4 donors).

**(C)** HIV-1 RNA levels in supernatants were quantified.

**(D)** CD69 and CD25 expression on CD4$^+$ T cells were analyzed at 72 hours (summary data shown).

Data in (C and D) are presented as mean ± SEM. Statistical significance was determined by paired Student's t-test (C and D). ns, no significance.

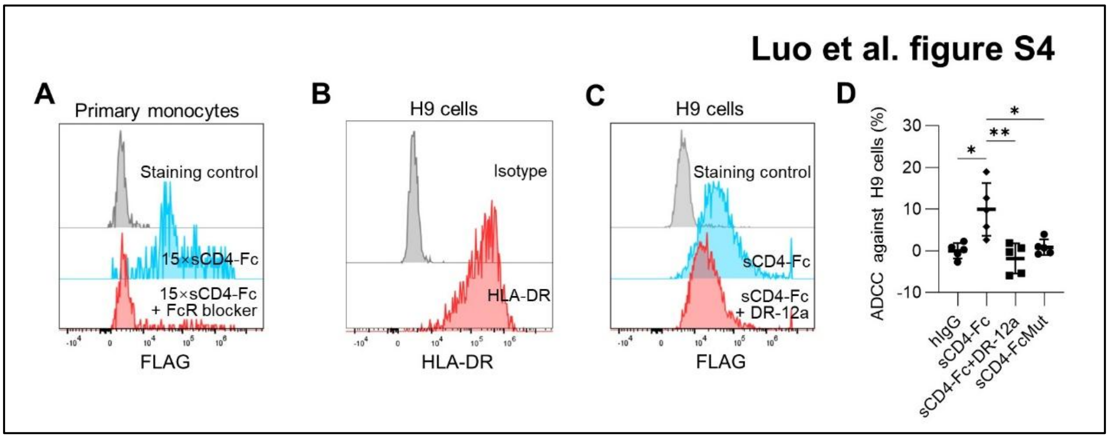


**Figure S4. sCD4-Fc mediates ADCC against HLA-DR$^+$ cells depending on HLA-DR**

**(A)** PBMCs from healthy donors were pre-treated with or without FcR blockade followed by incubation with 15×sCD4-Fc (20 nM) for 30 minutes and then staining with anti-FLAG and CD14. Representative flow histograms showed the biding of 15×sCD4-Fc to CD14$^+$ monocytes.

**(B)** Flow cytometric analysis of HLA-DR surface expression on H9 cells (histogram). Shaded: isotype control.

**(C)** HLA-DR-dependent binding blockade. H9 cells were incubated with sCD4-Fc (20 nM) with or without DR-12a peptides (50 µg/mL) for 3 hours at 37 °C. Binding was detected via anti-FLAG-APC staining (representative of 2 experiments).

**(D)** Mechanism of sCD4-mediated cytotoxicity. Primary human NK cells (effectors) were co-cultured with H9 target cells at a 30:1 E:T ratio for 6 hours under four experimental conditions: (i) hIgG1 control (100 nM), (ii) sCD4-Fc (100 nM), (iii) sCD4-Fc precomplexed with DR-12a blocking peptide (50 µg/mL, 20 min incubation at room temperature), or (iv) sCD4-FcMut (P329G/L234A/L235A; 100 nM). Viable H9 cells were quantified by flow cytometry, with specific lysis calculated (representative of 2 experiments).

Data in (D) are presented as mean ± SEM. Statistical significance was determined by was determined by one-way ANOVA followed by Tukey's multiple comparisons test (D). $*p < 0.05$, $**p < 0.01$.

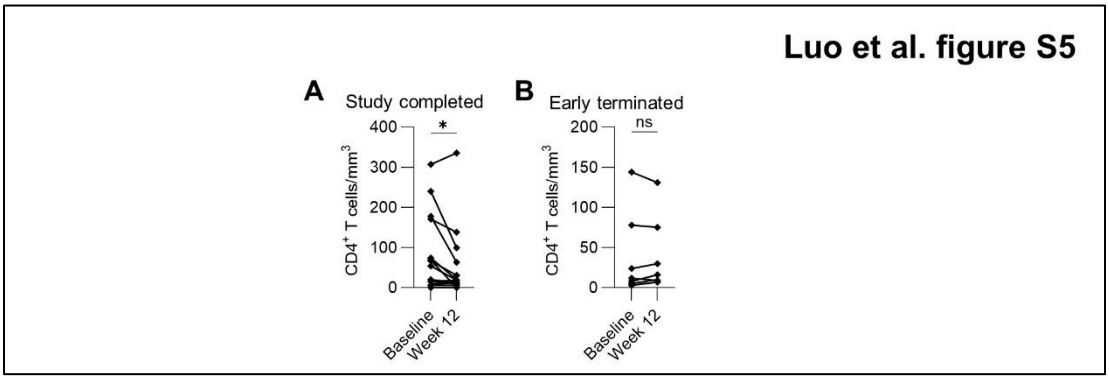


**Figure S5. Re-analysis of published clinical data indicates $CD4^+$ T-cell depletion following sCD4-Fc treatment in PLWH**

Re-analysis of published clinical data (Reference #52). Twenty-two subjects with advanced HIV-1 infection were enrolled in the study and treated intravenously with CD4-IgG (dosing details provided in the reference). Baseline $CD4^+$ T cell levels and week 12 counts were reanalyzed based on study completion. Summary data show changes in $CD4^+$ T cell counts for the 14 subjects who completed the 12-week treatment (A) versus the 8 participants who discontinued prematurely (B). Data are presented as paired data plot. Statistical significance was determined by paired Student's t-test (c). ns, no significance, $*p < 0.05$.

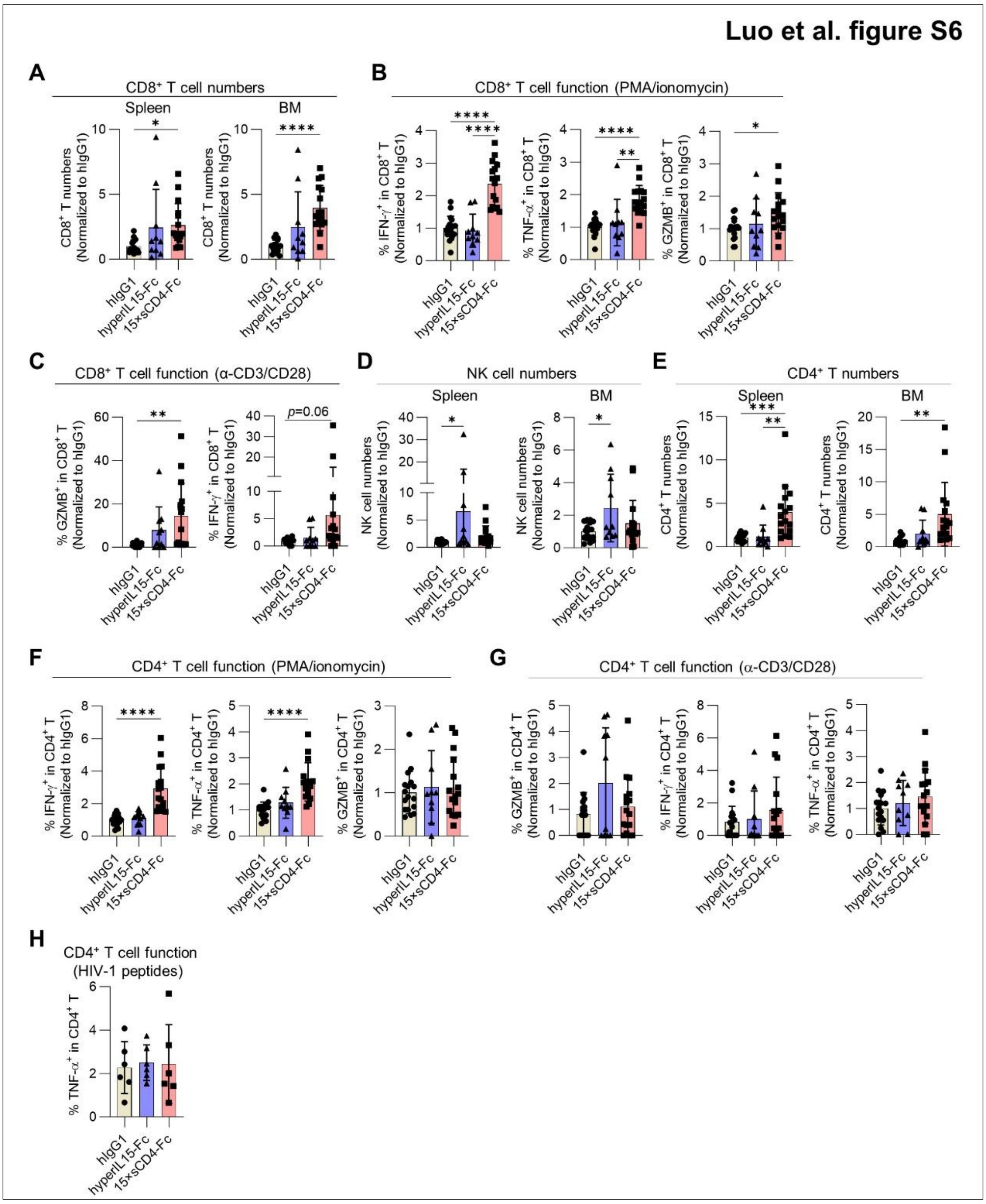


**Figure S6. The immunological effects of 15×sCD4-Fc treatment in cART-suppressed HIV-1-infected humanized mice**

Hu-mice were treated as in Figure 5. At the experimental endpoint, animals were euthanized, and lymphoid organs were harvested for analysis of human immune cell populations and functions.

**(A)** Relative $CD8^+$ T cell numbers in spleen and bone marrow. Data normalized to hIgG1 controls.

**(B)** $CD8^+$ T cell response to PMA/ionomycin stimulation. Splenocytes from hIgG1, hyperIL-15-Fc- or 15×sCD4-Fc- treated groups were stimulated with PMA (50 ng/mL) and ionomycin (1 μM) for 5 hours in the presence of brefeldin A, followed by flow cytometric detection of granzyme B (GZMB), IFN-γ, and TNF-α in $CD8^+$ T cells.

**(C)** Polyclonal $CD8^+$ T cell responses. Splenocytes were stimulated with α-CD3/CD28 for 8 hours (brefeldin A added at 3 hours). Intracellular granzyme B (GZMB) and IFN-γ in $CD8^+$ T cells were detected by flow cytometry.

**(D-E)** Relative NK (D) and $CD4^+$ T cell (E) numbers in spleen and bone marrow at endpoint. Data normalized to hIgG1 controls.

**(F-G)** $CD4^+$ T cell response to PMA/ionomycin or α-CD3/CD28 stimulation. Cytokine production in $CD4^+$ T cells after stimulation with PMA/ionomycin or (F) α-CD3/CD28 (G), measured as in (B-C).

**(H)** Antigen-specific $CD4^+$ T cell responses. Mixture of splenocytes and BM cells were stimulated ex vivo with HIV-1 peptide pools in the presence of α-CD28 and CD49d for 8 hours (Brefeldin A added at 3 hours), followed by intracellular TNF-α detection in $CD4^+$ T cells.

Data are presented as mean ± SEM. Statistical significance was determined by one-way ANOVA followed by Tukey's post hoc multiple comparisons test. $^*p < 0.05$, $^{**}p < 0.01$, $^{***}p < 0.001$, $^{****}p < 0.0001$.

### Table S1. Summary information about ART-treated patients (Related to Figure 1-3)

| | Sample ID. | Gender | Age | HIV-1 diagnosis date | The beginning date of ART | Sampling data | Plasma viral load at sampling | CD4⁺ T number at sampling (counts/μl) |
|---|---|---|---|---|---|---|---|---|
| Related to Figure 1F-I, Figure 3C-F and Figure S2C | BY155 | Male | 25 | 9/26/2021 | 9/30/2021 | 6/14/2025 | TND[A] | 1204 |
| | BY156 | Female | 38 | 10/25/2017 | 10/31/2018 | 6/14/2025 | TND | 513 |
| | BY157 | Male | 45 | 10/20/2005 | 10/25/2005 | 6/14/2025 | TND | 619 |
| | BY158 | Male | 44 | 10/12/2017 | 11/18/2018 | 6/14/2025 | TND | 666 |
| Related to Figure 1F-I, Figure 3C-D and Figure S2C | BY159 | Male | 70 | 12/14/2019 | 12/17/2019 | 6/14/2025 | TND | 519 |
| | BY160 | Male | 37 | 11/23/2017 | 9/5/2018 | 6/14/2025 | TND | 634 |
| | BY161 | Male | 41 | 6/12/2024 | 6/18/2024 | 6/14/2025 | TND | 543 |
| | BY162 | Male | 34 | 7/30/2017 | 9/11/2018 | 6/14/2025 | TND | 461 |
| Related to Figure 1J-K and Figure S2A-B, S2D-E | D29 | Male | 21 | 6/23/2021 | 10/22/2021 | 10/17/2022 | TND | 571 |
| | D30 | Male | 22 | 10/14/2020 | 10/22/2020 | 10/24/2022 | TND | 589 |
| | A1 | Male | 46 | 4/27/2012 | 7/29/2013 | 2/21/2023 | TND | 511 |
| | A2 | Male | 21 | 9/30/2021 | 11/14/2021 | 2/21/2023 | TND | 575 |
| | A3 | Male | 24 | 6/28/2017 | 11/7/2017 | 2/21/2023 | TND | 627 |
| | A4 | Male | 29 | 12/11/2018 | 11/22/2019 | 2/21/2023 | TND | 685 |
| | A5 | Male | 23 | 5/27/2020 | 6/16/2020 | 3/3/2023 | TND | 399 |
| | A6 | Male | 23 | 8/27/2020 | 9/12/2020 | 3/3/2023 | TND | 560 |
| Related to Figure 2B | BY169 | Female | 47 | 12/21/2010 | 1/3/2011 | 6/24/2025 | TND[A] | 632 |
| | BY170 | Male | 37 | 3/27/2012 | 4/4/2012 | 6/24/2025 | TND | 663 |
| | BY171 | Male | 50 | 12/4/2016 | 12/18/2016 | 6/24/2025 | TND | 537 |
| | BY172 | Male | 46 | 8/18/2019 | 8/25/2019 | 6/24/2025 | TND | 747 |
| | BY173 | Male | 46 | 9/22/2015 | 10/1/2015 | 6/24/2025 | TND | 713 |
| Related to Figure 2C-D | BY191 | Male | 57 | 12/8/2015 | 10/12/2018 | 7/4/2025 | TND | 594 |
| | BY192 | Male | 75 | 11/3/2016 | 11/6/2018 | 7/4/2025 | TND | 882 |
| | BY193 | Male | 60 | 11/8/2012 | 9/19/2018 | 7/4/2025 | TND | 911 |
| | BY195 | Male | 45 | 10/13/2016 | 10/25/2018 | 7/4/2025 | TND | 1151 |
| | BY197 | Male | 48 | 11/29/2018 | 12/5/2018 | 7/4/2025 | TND | 594 |
| Related to Figure 2O-Q | BY152 | Male | 36 | 9/16/2014 | 9/22/2017 | 5/30/2025 | TND | 1005 |
| | BY153 | Male | 33 | 9/8/2015 | 9/24/2015 | 5/30/2025 | TND | 817 |
| Related to Figure 3D-F | BY131 | Male | 68 | 7/11/2019 | 8/22/2019 | 5/24/2025 | TND | 625 |
| | BY133 | Male | 37 | 6/27/2018 | 8/27/2018 | 5/24/2025 | TND | 584 |
| | BY134 | Male | 35 | 6/22/2019 | 6/30/2019 | 5/24/2025 | TND | 492 |
| | BY137 | Male | 45 | 4/19/2007 | 5/26/2007 | 5/24/2025 | TND | 710 |
| Related to Figure 3D | 128 | Male | 34 | 10/16/2013 | 10/1/2023 | 5/29/2025 | TND | 1080 |
| | 130 | Male | 25 | 7/25/2023 | 9/7/2023 | 5/30/2025 | TND | 692 |
| | 132 | Male | 60 | 9/21/2016 | 10/8/2016 | 5/30/2025 | TND | 791 |
| Related to Figure 3E-F | BY130 | Male | 31 | 7/10/2022 | 7/14/2022 | 5/24/2025 | TND | 817 |
| | BY132 | Male | 42 | 7/26/2018 | 10/11/2018 | 5/24/2025 | TND | 524 |
| | BY135 | Male | 35 | 6/2/2016 | 10/30/2016 | 5/24/2025 | TND | 557 |

| | | | | | | | | |
|---|---|---|---|---|---|---|---|---|
| Related to Figure S3C-D | BY187 | Male | 26 | 3/3/2022 | 3/10/2022 | 7/3/2025 | TND | 902 |
| | BY188 | Male | 23 | 8/13/2017 | 9/15/2017 | 7/3/2025 | TND | 845 |
| | BY189 | Male | 28 | 11/6/2018 | 12/1/2018 | 7/3/2025 | TND | 670 |
| | BY190 | Male | 65 | 12/1/2002 | 12/10/2002 | 7/3/2025 | TND | 884 |

[A]TND: Target not detected (limit of detection <20 copies/mL)

**Table S2. sCD4-Fc treatment induces systemic human lymphocytes depletion in HIV-1-infected humanized mice (Related to Figure 4)**

| Cohort | Group | Mouse # | Pre-treatment hCD45 (W7) | HIV-1 RNA in plasma (log10 IU/mL)[B] | | | | | Number of splenic human immune subsets (W9.5) | | | | Cell-associated HIV-1 DNA[C] | |
|---|---|---|---|---|---|---|---|---|---|---|---|---|---|---|
| | | | hCD45 (%)[A] | W3 | W7 | W7.35 | W8 | W9.5 | hCD45 (×10$^6$) | CD8$^+$ T (×10$^4$) | CD4$^+$ T (×10$^4$) | NK (×10$^4$) | SP | BM |
| 1 | hIgG1 | 442 | 24.60 | 5.47 | TND[D] | TND | TND | TND | 1.29 | 40.90 | 20.60 | 3.06 | N.A[E] | N.A |
| | | 446 | 70.80 | 4.86 | 2.51 | 2.86 | 2.7 | TND | 2.91 | 66.97 | 77.69 | 6.52 | N.A | N.A |
| | | 450 | 69.4 | 4.80 | TND | TND | TND | TND | 4.32 | 103.13 | 93.67 | 8.51 | N.A | N.A |
| | | 471 | 77.4 | 4.87 | TND | TND | TND | TND | 3.72 | 97.85 | 67.04 | 11.01 | N.A | N.A |
| | sCD4-Fc | 464 | 44.92 | 4.56 | TND | TND | TND | TND | 0.92 | 20.13 | 4.35 | 2.84 | N.A | N.A |
| | | 449 | 55.01 | 4.51 | 2.61 | TND | TND | TND | 0.17 | 4.03 | 0.97 | 0.37 | N.A | N.A |
| | | 461 | 66.00 | 4.60 | TND | TND | TND | TND | 0.30 | 9.18 | 1.62 | 0.86 | N.A | N.A |
| | | 469 | 72.93 | 4.80 | TND | TND | TND | TND | 0.77 | 15.49 | 3.32 | 2.56 | N.A | N.A |
| | | 456 | 73.38 | 4.59 | TND | TND | TND | TND | 1.26 | 26.58 | 10.39 | 2.23 | N.A | N.A |
| | 15× sCD4-Fc | 449 | 51.12 | 4.10 | TND | TND | TND | TND | 5.37 | 152.00 | 71.50 | 20.80 | N.A | N.A |
| | | 458 | 70.73 | 4.64 | TND | TND | 2.42 | TND | 10.30 | 350.00 | 247.00 | 25.30 | N.A | N.A |
| | | 455 | 62.01 | 4.32 | TND | TND | TND | TND | 10.60 | 151.00 | 420.00 | 12.10 | N.A | N.A |
| | | 466 | 54.19 | 4.87 | TND | TND | TND | TND | 6.12 | 205.00 | 162.00 | 10.20 | N.A | N.A |
| 2 | hIgG1 | 031 | 58.45 | 4.33 | TND | TND | TND | TND | 20.30 | 402.73 | 216.14 | 81.60 | 1.02 | 0.94 |
| | | 037 | 62.84 | 4.56 | TND | TND | TND | TND | 24.19 | 568.49 | 153.72 | 80.79 | 0.94 | 1.74 |
| | | 038 | 64.23 | 4.49 | TND | TND | TND | TND | 34.11 | 577.08 | 214.48 | 85.95 | 0.65 | 0.28 |
| | | 047 | 65.17 | 4.51 | TND | TND | TND | TND | 24.65 | 402.97 | 218.95 | 114.86 | 2.06 | 1.19 |
| | | 050 | 65.48 | 3.87 | TND | TND | TND | TND | 25.18 | 565.53 | 221.99 | 83.33 | 0.33 | 0.85 |
| | sCD4-Fc | 033 | 81.51 | 3.88 | TND | TND | TND | TND | 6.08 | 115.91 | 35.79 | 12.34 | 0.52 | 0.94 |
| | | 045 | 73.96 | 4.12 | TND | TND | TND | TND | 8.16 | 120.88 | 33.29 | 18.84 | 1.71 | 1.74 |
| | | 051 | 27.42 | 2.71 | TND | TND | TND | TND | 6.92 | 91.62 | 37.55 | 16.46 | 0.34 | 0.28 |
| | sCD4-FcMut | 035 | 45.10 | 4.23 | TND | TND | TND | TND | 13.29 | 374.46 | 80.36 | 56.10 | 0.60 | 0.67 |
| | | 042 | 54.86 | 4.70 | TND | TND | TND | TND | 22.05 | 460.17 | 198.47 | 117.51 | 0.49 | 0.93 |
| | | 048 | 58.05 | 4.49 | TND | TND | TND | TND | 27.04 | 700.23 | 236.81 | 82.75 | 1.07 | 1.81 |
| | | 049 | 86.13 | 4.01 | TND | TND | TND | TND | 18.31 | 255.66 | 166.31 | 51.44 | 1.12 | 0.72 |
| | | 052 | 79.18 | 3.78 | TND | TND | TND | TND | 25.39 | 724.88 | 224.45 | 117.05 | 0.34 | 0.70 |
| 3 | hIgG1 | 66 | 92.91 | 4.32 | TND | TND | TND | TND | 33.79 | 559.75 | 2170.80 | 63.52 | 0.29 | 0.70 |
| | | 111 | 49.06 | 4.62 | 3.33 | TND | TND | TND | 12.84 | 257.29 | 620.89 | 60.58 | 0.45 | 0.65 |
| | | 492 | 49.06 | 4.86 | TND | TND | TND | TND | 28.94 | 413.74 | 830.09 | 196.21 | 2.25 | 1.65 |
| | sCD4-FcMut | 060 | 72.76 | 4.15 | 2.49 | TND | TND | TND | 12.86 | 192.96 | 592.07 | 80.14 | 0.57 | 1.10 |
| | | 114 | 64.51 | 4.49 | 3.19 | TND | TND | TND | 10.09 | 153.03 | 333.97 | 71.64 | 1.31 | 2.53 |
| | | 465 | 29.13 | 5.08 | 3.22 | 2.75 | TND | TND | 7.32 | 170.89 | 441.67 | 34.85 | 0.51 | 2.34 |

Humanized mice were treated as described in Figure 4. Cohorts 1-3 consisted of NCG-hu HSC mice. [A]Percentage of hCD45 in peripheral blood at week 7. [B]Plasma viral load at indicated weeks post HIV-1 infection. [C]Relative levels, normalized to hIgG1 grop. [D]TND: Target not detected (detection limit <250 IU/mL). [E]N.A: No Applicable.

**Table S3. 15×sCD4-Fc treatment reactivates HIV-1 reservoir, enhances immune functions and decreases HIV reservoir size in HIV-1-infected humanized mice (Related to Figure 5)**

| Cohort | Group | Mouse # | Pre-treatment hCD45 (W7) | HIV-1 RNA in plasma (log10 IU/mL)[B] | | | | | Number of splenic human immune subsets | | | | Cell-associated HIV-1 DNA[C] | | IPDA[D] | | TZA[E] |
|---|---|---|---|---|---|---|---|---|---|---|---|---|---|---|---|---|---|
| | | | hCD 45 (%)[A] | W3 | W7 | W 7.35 | W9 | W 11.5 | hCD 45 $(10^6)$ | $CD8^+$ T $(10^4)$ | $CD4^+$ T (104) | NK $(10^4)$ | SP | BM | SP | BM | BM |
| 1 | hIgG1 | 442 | 24.62 | 5.47 | TND[F] | TND | N.A[G] | N.A | 1.29 | 40.9 | 20.6 | 3.06 | N.A | N.A | N.A | N.A | N.A |
| | | 446 | 70.81 | 4.86 | 2.51 | 2.86 | N.A | N.A | 2.91 | 67.0 | 62.5 | 6.52 | N.A | N.A | N.A | N.A | N.A |
| | | 450 | 69.44 | 4.80 | TND | TND | N.A | N.A | 4.32 | 103.0 | 93.7 | 8.51 | N.A | N.A | N.A | N.A | 137.1 |
| | | 471 | 77.47 | 4.87 | TND | TND | N.A | N.A | 3.72 | 97.9 | 67.0 | 11.0 | N.A | N.A | N.A | N.A | 287.1 |
| | 15× sCD4-Fc | 449 | 51.12 | 4.10 | TND | TND | N.A | N.A | 5.37 | 152.0 | 71.5 | 20.8 | N.A | N.A | N.A | N.A | 37.0 |
| | | 458 | 70.73 | 4.64 | TND | TND | N.A | N.A | 10.30 | 350.0 | 247.0 | 25.3 | N.A | N.A | N.A | N.A | 0.00 |
| | | 455 | 62.01 | 4.32 | TND | TND | N.A | N.A | 10.60 | 151.0 | 420.0 | 12.1 | N.A | N.A | N.A | N.A | 34.5 |
| | | 466 | 54.19 | 4.87 | TND | TND | N.A | N.A | 6.12 | 205.0 | 162.0 | 10.2 | N.A | N.A | N.A | N.A | 18.3 |
| 4 | hIgG1 | 308 | 86.65 | 4.10 | TND | TND | TND | N.A | 35.40 | 402.6 | 120.6 | 424.8 | 0.47 | 0.58 | 0.00 | 0.20 | N.A |
| | | 313 | 72.91 | 4.29 | TND | TND | TND | N.A | 29.60 | 197.8 | 142.5 | 293.0 | 0.34 | 0.26 | 0.50 | 0.25 | N.A |
| | | 317 | 82.33 | 4.66 | TND | TND | TND | N.A | 18.92 | 194.3 | 84.9 | 208.1 | 2.19 | 2.16 | 2.50 | 2.54 | N.A |
| | 15× sCD4-Fc | 309 | 54.55 | 3.93 | TND | TND | TND | N.A | 30.00 | 238.9 | 113.7 | 360.0 | 0.53 | 0.27 | 0.00 | 0.10 | N.A |
| | | 312 | 82.25 | 4.67 | TND | TND | TND | N.A | 19.12 | 254.2 | 132.1 | 286.8 | 1.08 | 0.55 | 1.21 | 0.54 | N.A |
| | | 315 | 85.95 | 4.14 | TND | TND | TND | N.A | 25.80 | 230.8 | 516.5 | 309.6 | 0.19 | 0.09 | 0.23 | 0.00 | N.A |
| 5 | hIgG1 | 058 | 74.62 | 4.98 | TND | TND | TND | TND | 3.26 | 49.8 | 45.8 | 16.6 | 0.92 | 0.77 | 0.90 | 1.39 | 501.7 |
| | | 059 | 74.25 | 4.92 | 2.51 | TND | TND | TND | 2.45 | 39.2 | 41.3 | 10.8 | 1.36 | 2.34 | 1.02 | 1.43 | 37.6 |
| | | 061 | 78.47 | 4.44 | TND | TND | TND | TND | 9.20 | 208.0 | 110.0 | 29.2 | 0.77 | 0.40 | 1.19 | 0.47 | 192.8 |
| | | 077 | 50.30 | 4.49 | TND | TND | TND | TND | 3.66 | 84.1 | 84.0 | 21.9 | 0.95 | 0.49 | 0.90 | 0.71 | 649.0 |
| | hyperIL-15-Fc | 060 | 75.67 | 4.33 | TND | TND | TND | TND | 10.60 | 69.4 | 37.8 | 635.0 | 1.23 | 0.33 | 3.41 | 0.08 | 108.4 |
| | | 067 | 76.21 | 4.76 | 2.47 | TND | TND | TND | 8.00 | 246.0 | 50.6 | 300.0 | 1.12 | 0.53 | 1.27 | 1.16 | 603.9 |
| | | 070 | 44.31 | 3.84 | TND | TND | TND | TND | 7.20 | 130.0 | 62.6 | 40.0 | 2.40 | 1.51 | 0.45 | 0.60 | 337.2 |
| | | 075 | 51.38 | 4.27 | TND | TND | TND | TND | 4.88 | 133.0 | 65.8 | 23.5 | 0.56 | 0.43 | 0.21 | 0.88 | 89.8 |
| | 15×sCD4-Fc | 063 | 62.42 | 4.71 | TND | TND | TND | TND | 4.56 | 154.0 | 180.4 | 22.2 | 0.42 | 0.31 | 0.58 | 0.65 | 160.1 |
| | | 064 | 75.76 | 4.42 | 2.74 | TND | 2.59 | TND | 5.84 | 140.0 | 127.00 | 39.3 | 0.69 | 0.36 | 1.66 | 0.34 | 0.00 |
| | | 069 | 57.76 | 4.70 | TND | TND | TND | TND | 12.0 | 319.0 | 429.0 | 32.7 | 1.10 | 0.16 | 0.39 | 0.27 | 49.8 |
| 6 | hIgG1 | 266 | 62.72 | 4.60 | TND | TND | TND | TND | 3.48 | 33.9 | 28.3 | 14.9 | 0.76 | 0.78 | 0.71 | 0.00 | 39.8 |
| | | 253 | 45.19 | 4.91 | TND | TND | TND | TND | 1.72 | 24.0 | 77.7 | 9.92 | 1.24 | 1.22 | 1.29 | 2.00 | 475.27 |
| | | 258 | 20.10 | 4.74 | TND | TND | TND | TND | 4.50 | 77.8 | 24.2 | 21.9 | N.A | N.A | N.A | N.A | N.A |
| | Hyper IL-15-Fc | 262 | 43.56 | 3.97 | TND | TND | TND | TND | 3.66 | 32.6 | 13.4 | 13.2 | 0.38 | 0.21 | 3.38 | 2.61 | 615.2 |
| | | 267 | 60.18 | 4.19 | TND | TND | TND | TND | 2.15 | 18.6 | 12.1 | 8.34 | 0.19 | 1.16 | 2.56 | 2.16 | 820.5 |
| | | 268 | 46.23 | 4.70 | TND | TND | TND | TND | 4.75 | 74.8 | 41.7 | 19.0 | N.A | N.A | N.A | N.A | N.A |
| | 15× sCD4-Fc | 261 | 55.08 | 4.42 | TND | 2.58 | TND | TND | 12.00 | 251.0 | 244.0 | 115.0 | 0.54 | 0.27 | 0.88 | 0.03 | 0.00 |
| | | 264 | 51.02 | 4.44 | TND | TND | TND | TND | 6.80 | 161.0 | 202.0 | 76.2 | 0.50 | 1.21 | 0.29 | 0.00 | 30.4 |
| | | 269 | 33.48 | 4.23 | TND | TND | TND | TND | 5.70 | 80.2 | 75.7 | 27.2 | N.A | N.A | N.A | N.A | N.A |

| | | | | | | | | | | | | | | | | | |
|---|---|---|---|---|---|---|---|---|---|---|---|---|---|---|---|---|---|
| 7 | hIgG1 | 105 | 55.88 | 4.27 | TND | TND | N.A | N.A | 16.60 | 686.0 | 453.0 | 106.0 | 1.46 | 1.60 | 0.36 | 0.56 | 10.4 |
| | | 113 | 29.94 | 4.25 | TND | 2.51 | N.A | N.A | 9.60 | 372.0 | 195.0 | 105.0 | 0.69 | 0.40 | 1.89 | 2.23 | 64.1 |
| | | 118 | 83.04 | 4.35 | TND | 2.57 | N.A | N.A | N.A | N.A | N.A | N.A | N.A | N.A | N.A | N.A | N.A |
| | | 127 | 29.05 | 3.95 | TND | TND | N.A | N.A | N.A | N.A | N.A | N.A | N.A | N.A | N.A | N.A | N.A |
| | | 131 | 33.41 | 4.33 | TND | TND | N.A | N.A | 9.90 | 333.0 | 332.0 | 122.0 | N.A | N.A | N.A | N.A | N.A |
| | hyperIL-15-Fc | 111 | 34.49 | 4.37 | TND | 2.48 | N.A | N.A | 65.30 | 4360.0 | 670.0 | 862.0 | 1.56 | 0.86 | 1.71 | 0.17 | 23.3 |
| | | 115 | 60.02 | 4.91 | TND | TND | N.A | N.A | 53.00 | 2720.0 | 1510.0 | 374.0 | 0.89 | 0.80 | 1.09 | 0.07 | 19.4 |
| | | 119 | 60.15 | 4.40 | TND | TND | N.A | N.A | N.A | N.A | N.A | N.A | N.A | N.A | N.A | N.A | N.A |
| | | 123 | 23.14 | 4.16 | 2.46 | 2.58 | N.A | N.A | 4.46 | 73.3 | 8.09 | 70.9 | N.A | N.A | N.A | N.A | N.A |
| | | 130 | 49.37 | 4.36 | TND | TND | N.A | N.A | N.A | N.A | N.A | N.A | N.A | N.A | N.A | N.A | N.A |
| | 15×sCD4-Fc | 101 | 45.16 | 3.37 | TND | 2.90 | N.A | N.A | 85.50 | 3050.0 | 4240.0 | 190.0 | 0.19 | 0.39 | 0.42 | 0.07 | N.A |
| | | 103 | 61.05 | 3.87 | TND | 2.75 | N.A | N.A | N.A | N.A | N.A | N.A | N.A | N.A | N.A | N.A | N.A |
| | | 122 | 40.24 | 4.51 | TND | 2.43 | N.A | N.A | 22.00 | 927.0 | 957.0 | 29.7 | 0.49 | 0.23 | 0.62 | 0.09 | 0.00 |
| | | 128 | 38.62 | 3.67 | TND | 2.52 | N.A | N.A | 30.00 | N.A | N.A | N.A | 0.49 | 0.14 | 0.75 | 0.00 | N.A |
| | | 129 | 27.31 | 4.12 | TND | 3.17 | N.A | N.A | N.A | 855.0 | 1160.0 | 98.4 | N.A | N.A | N.A | N.A | N.A |
| | | 134 | 45.33 | 3.58 | TND | 2.83 | N.A | N.A | N.A | N.A | N.A | N.A | N.A | N.A | N.A | N.A | N.A |

Humanized mice were treated as described in Figure 5. Cohorts 1 and 4-6 consisted of NCG-hu HSC mice, while cohort 7 comprised NCG-hu Thy/HSC mice. [A]Percentage of hCD45 in peripheral blood at week 7. [B]Plasma viral load at indicated weeks post HIV-1 infection. [C]Relative levels, normalized to hIgG1 group. [D]Intact provirus DNA (relative levels, normalized to hIgG1 grop). [E]TZM-bl based quantitative assay. [F]TND: Target not detected (detection limit <250 IU/mL). [G]N.A: No Applicable.
